# Proteome profiling identifies a link between the mitochondrial pathways and host-microbial sensor ELMO1 following *Salmonella* infection

**DOI:** 10.1101/2024.05.03.592405

**Authors:** Sajan C Achi, Dominic McGrosso, Stefania Tocci, Stella-Rita Ibeawuchi, Ibrahim M. Sayed, David J Gonzalez, Soumita Das

## Abstract

The host EnguLfment and cell MOtility protein 1 (ELMO1) is a cytosolic microbial sensor that facilitates bacterial sensing, internalization, clearance, and inflammatory responses. We have shown previously that ELMO1 binds bacterial effector proteins, including pathogenic effectors from *Salmonella* and controls host innate immune signaling. To understand the ELMO1-regulated host pathways, we have performed liquid chromatography Multinotch MS3-Tandem Mass Tag (TMT) multiplexed proteomics to determine the global quantification of proteins regulated by ELMO1 in macrophages during *Salmonella* infection. Comparative proteome analysis of control and ELMO1-depleted murine J774 macrophages after *Salmonella* infection quantified more than 7000 proteins with a notable enrichment in mitochondrial-related proteins. Gene ontology enrichment analysis revealed 19 upregulated and 11 downregulated proteins exclusive to ELMO1-depleted cells during infection, belonging to mitochondrial functions, metabolism, vesicle transport, and the immune system. By assessing the cellular energetics via Seahorse analysis, we found that *Salmonella* infection alters mitochondrial metabolism, shifting it from oxidative phosphorylation to glycolysis. Importantly, these metabolic changes are significantly influenced by the depletion of ELMO1. Furthermore, ELMO1 depletion resulted in a decreased ATP rate index following *Salmonella* infection, indicating its importance in counteracting the effects of *Salmonella* on immunometabolism. Among the proteins involved in mitochondrial pathways, mitochondrial fission protein DRP1 was significantly upregulated in ELMO1-depleted cells and in ELMO1-KO mice intestine following *Salmonella* infection. Pharmacological Inhibition of DRP1 revealed the link of the ELMO1-DRP1 pathway in regulating the pro-inflammatory cytokine TNF-α following infection. The role of ELMO1 has been further characterized by a proteome profile of ELMO1-depleted macrophage infected with SifA mutant and showed the involvement of ELMO1-SifA on mitochondrial function, metabolism and host immune/defense responses. Collectively, these findings unveil a novel role for ELMO1 in modulating mitochondrial functions, potentially pivotal in modulating inflammatory responses.

**Graphical Abstract:** 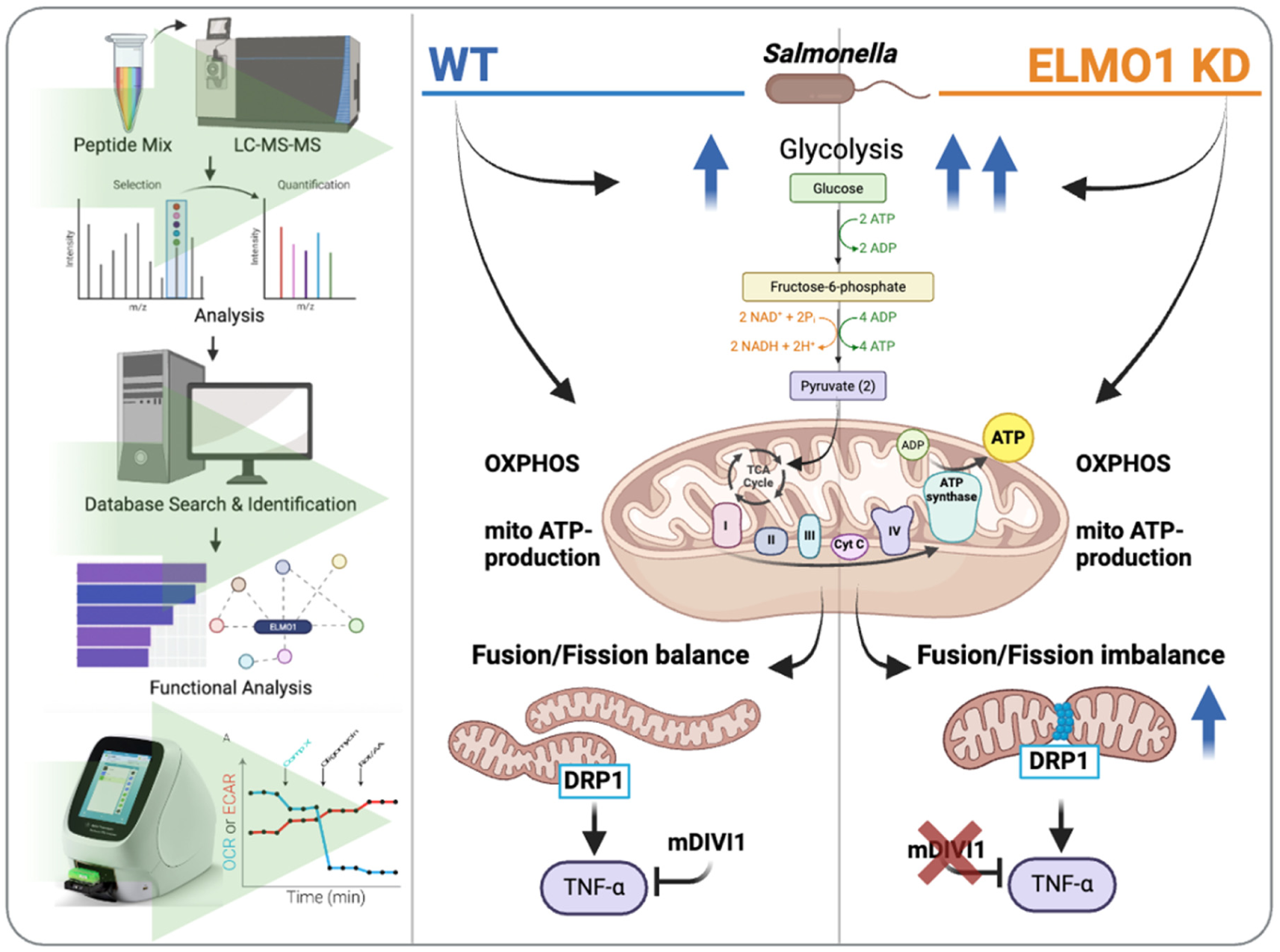

**Significance Statement:** Host microbial sensing is critical in infection and inflammation. Among these sensors, ELMO1 has emerged as a key regulator, finely tuning innate immune signaling and discriminating between pathogenic and non-pathogenic bacteria through interactions with microbial effectors like SifA of *Salmonella*. In this study, we employed Multinotch MS3-Tandem Mass Tag (TMT) multiplexed proteomics to determine the proteome alterations mediated by ELMO1 in macrophages following WT and SifA mutant *Salmonella* infection. Our findings highlight a substantial enrichment of host proteins associated with metabolic pathways and mitochondrial functions. Notably, we validated the mitochondrial fission protein DRP1 that is upregulated in ELMO1-depleted macrophages and in ELMO1 knockout mice intestine after infection. Furthermore, we demonstrated that *Salmonella*-induced changes in cellular energetics are influenced by the presence of ELMO1. This work shed light on a possible novel link between mitochondrial dynamics and microbial sensing in modulating immune responses.

## Introduction

Host cellular responses coordinate during infections to maintain immune homeostasis. Infection with enteric pathogens generates inflammation and tissue damage which leads to 4-6 million human deaths worldwide per year. Microbial sensing controls inflammation after recognizing invading pathogens and regulates the expression of proteins critical for the disease outcome. Understanding the molecular mechanisms governing the host-pathogen interaction is vital for controlling and treating infectious diseases worldwide. To this end, mass spectrometry (MS) has become a powerful and effective approach in proteomics to better understand complex and dynamic host-pathogen interactions in unbiased fashions. Recent breakthroughs in the proteomics field have included the use of isobaric labels termed Tandem Mass Tags (TMT). The TMT approach has been shown to yield data of high quantitative accuracy, when LC-MS3 level quantification using the synchronous precursor selection (SPS) is performed. *Salmonella* has been used as a model enteric bacterial pathogen and the proteome profile of the bacteria has been extensively studied (1–4). In contrast, how the host proteome fluctuates during infection is less understood given the complexity of the host proteome and the number of proteins that present with greater dynamic ranges. Previous studies using LCMS/MS of RAW 264.7 macrophages after *Salmonella* infection identified the involvement of proteins in the production of antibacterial nitric oxide, production of prostaglandin H2, and regulation of intracellular traffic (5). A global host phosphoproteome analysis after bacterial infection identified the kinase SIK2 as a central component of the host defense machinery upon *Salmonella* infection(6).

Though host transcriptome profiling has been widely practiced in host-pathogen interactions (7), only a few studies reported multiplex proteomic profiling of host proteome upon infection. Proteomics has been used to study host protein profiles during infection, either studying the whole cell proteome (5, 8) or subsets of the host proteome such as Golgi networks (9, 10) and *Salmonella*-containing vacuoles (SCVs)(11). To understand the host cellular pathways modified in macrophages after *Salmonella* infection, here, we have used tandem mass tags, where chemical labeling with combined samples reduces errors associated with run-to-run variations caused by temperature or column conditions (12).

For microbial sensing Pattern Recognition Receptors (PRRs) recognize pathogen-associated molecular patterns (PAMPs) or host-derived damage-associated molecular patterns (DAMPs) and activate inflammatory signals (13–15). Our published work identified <u>B</u>rain <u>A</u>ngiogenesis Inhibitor1 (BAI1) as a new PRR which was initially identified as a receptor for apoptotic cells(16). The intracellular domain of BAI1 interacts with ELMO1 (<u>E</u>ngulfment and cel<u>l</u> <u>mo</u>tility protein 1), which associates with Dock180 (<u>D</u>edicator <u>o</u>f <u>c</u>yto<u>k</u>inesis 180) to act as a bipartite guanine nucleotide exchange factor (GEF) for the small GTPase Rac1 (16) and the activated Rac1 facilitates the engulfment of bacteria. We have shown that ELMO1 is more than just a director of phagocytosis (17) and involved in the internalization of enteric bacteria and generation of intestinal inflammation (18); ELMO1 is elevated in patients with inflammatory bowel disease and induces inflammatory cytokines (19, 20); ELMO1 regulates autophagy induction and bacterial clearance during infection with *Salmonella* (21); ELMO1 interacts with microbial sensor NOD2 and together with mutant NOD2 associated with Crohn’s disease (CD) are unable to clear CD-associated bacteria (22). Interestingly, ELMO1 differentially regulates the immune response after sensing pathogens and interacts with the bacterial effectors containing a signature motif called WxxxE, which is absent in the commensals (23, 24). The same work identified the *Salmonella* WxxxE-containing effector SifA interacts with ELMO1, and controls innate immune responses.

Here, we have used TMT-guided LC-MS3 proteomics to determine the global quantification of proteins regulated by ELMO1 in macrophages during *Salmonella* infection. Comparative proteome analysis of control and ELMO1-depleted murine macrophages (J774) after *Salmonella* (*SL*) infection quantified more than 7000 proteins and identified the mitochondrial pathway as a top candidate regulated by ELMO1. We also identified the differential proteome profile after infection of ELMO1-depleted macrophages with WT *SL* and *SifA* mutant *Salmonella*. Together, we report an extensive comparative proteome profile of ELMO1-regulated host proteome components following *Salmonella* infection.

## Materials and Methods

### Bacterial strains and growth conditions

*Salmonella enterica* serovar Typhimurium (strain *SL1344, abbreviated as SL*) and *sifA* mutant strain were used in this study following the culture method as described before (18, 24).

### Cell culture and infection

Culturing and maintenance of Control (C1) and ELMO1-depleted (E1) J774 macrophages (J774) were described before (18, 24). For infection, approximately 8 × 10^6^ cells were plated onto a 100 mm dish and cells were infected with SL at a multiplicity of infection (moi) of 1:10 for 1h. Extracellular bacteria were killed by gentamicin treatment (500 µg/mL) for 90 minutes at 37◦C, followed by treatment with low-dose gentamicin (50 µg/mL) for 3.5 h. Cell pellets were collected and stored at -80℃ to until processing.

### Preparation of samples for multiplex proteomics

Samples were lysed using a Q500 QSonica sonicator (Qsonica) equipped with 1.6 mm microtip at 20% amplitude in 200 µL of lysis buffer (6M urea, 7% SDS, 50 mM TEAB, pH 8.1 and complete protease inhibitor and PhosStop tablets [Roche]). Samples were reduced using 500 mM DTT at 47°C for 30 minutes and alkylated using 500 mM Iodoacetamide (IAA) in the dark at room temperature (RT) for, 45 minutes. After that, quenching was performed using 500 mM DTT at RT for 5 minutes. Each sample was then acidified using 12% phosphoric acid and mixed in a 7:1 ratio with binding buffer (90% Methanol, 50 mM TEAB, pH 7.1) and loaded into S-Trap columns (Protifi). On-column digestion occurred using 5 ug trypsin (Promega sequence-grade trypsin) in 50 mM TEAB for 3 hours at 47C. Peptides were sequentially eluted in 50 mM TEAB, 5% formic acid, and 50% acetonitrile 5% formic acid, and subsequently dried in a speed-vac. Samples were then desalted with C18 Sep-Paks (Waters) and eluted with a 40% and 80% acetonitrile solution containing 0.5% acetic acid. The peptide concentration was determined by the protein estimation kit (Pierce, Catalog # 23275), and 50 μg aliquots of each sample were dried in a speed-vac. To control for run-to-run variation, 10 µg of each sample was also pooled for “bridge” channels as previously described (25). Each sample was resuspended in 30% dry acetonitrile in 200 mM HEPES (pH 8.5) for Tandem Mass Tag labelling with 7 μl of the appropriate TMT reagent, as previously described (26). Labelling was performed at RT for 1 h and quenched with 8 μl 5% hydroxylamine (Sigma). Labelled samples were acidified by adding 50 μl 1% TFA. After TMT labelled, each 10-plex experiment was combined, desalted (Waters C18 Sep-Paks), and dried in a speed-vac. TMT manufacturer’s channel contamination was controlled according to using reported contamination values as recorded by the lot number (VL312003). Combined and labelled samples were subjected to basic pH reverse-phase liquid chromatography (LC, Thermo Scientific Ultimate 3000, Biobasic c18 column), collected in fractions as previously described (27), and dried to completion. Samples were then resuspended in 20 µL 5% formic acid, and 5% acetonitrile and placed in glass vials for future analysis.

### LC-MS^3^ strategies

Sample data was acquired using LC-MS/MS/MS using an Orbitrap Fusion mass spectrometer (Thermo Scientific) with an in-line EASY-nLC 1000 instrument (Thermo Scientific). Separation and acquisition settings were performed using previously defined methods(27).

Briefly, the fractions were run on three-hour gradients starting at 3% acetonitrile and 0.125% formic acid and ending at 100% acetonitrile and 0.125% formic acid. Peptides were separated on an in-house packed 30 cm × 100 μm inner diameter, 360 μm outer diameter column comprised of 0.5 cm C4 resin (diameter = 5 μm), 0.5 cm C18 resin (diameter = 3 μm), and 29 cm C18 resin (diameter = 1.8 μm). Source ionization was performed by applying 2000V of electricity through the T-junction joining sample, waste, and column capillary termini.

MS1 spectrum acquisition was performed in data-dependent mode with an Orbitrap survey scan range of 500 – 1200 m/z and a resolution of 60,000. Automatic gain control (AGC) 2×10^5^ with a maximum ion inject time of 100 ms. Top N was used with N = 10 for both MS2 and MS3 fragment ion analysis.

MS2 data were collected using the decision tree option. Ions carrying 2 charges were analyzed between 600 – 1200 m/z, and ions carrying 3 or 4 charges were analyzed between 500 – 1200 m/z. The ion intensity threshold was 5×10^4^. Selected ions were isolated in the quadrupole at 0.5 Th and fragmented via collision-induced dissociation (CID). Fragment ion detection and data centroiding occurred in the linear ion trap with a rapid scan rate AGC target of 1×10^4^.

TMT-based quantitation via MS3 fragmentation was performed using synchronous precursor selection. Up to 10 MS2 precursors were concurrently fragmented using High Energy Collisional Dissociation (HCD) fragmentation. Reporter ions were detected in the Orbitrap at a resolution of 60,000 and with a lower threshold of 110 m/z. AGC was set to 1×10^5 with a maximum ion inject time of 100 ms. Data collected was centroided and precursor ions outside of 40 m/z below and 15 m/z above the MS2 m/z were removed.

### Proteomic Data Analysis

Mass spectrometry data was searched using Proteome Discoverer 2.5 software (Thermo Fisher Scientific). Data was searched against the reference proteome for *Mus musculus* downloaded from Uniprot.com on 10/01/2021. The SEQUEST HT search algorithm was employed to align MS2 spectral data against theoretical peptides (28). Precursor tolerance was set to 50 ppm and fragment tolerance was set to 0.6 Da. Static modifications were specified for TMT labels on N-termini and lysine residues, as well as for carbamidomethylation of cysteines. Dynamic modifications were set for the oxidation of methionine. A 1% false discovery rate cutoff was specified for the decoy database search(29). After filtering for quality (S/N > 10, Isolation Interference < 25), we summed the remaining peptide spectral match abundances to the protein level, and resultant values were normalized against the average value for each protein divided by the median of all average protein values. A second normalization step was performed whereby the abundance value for each protein per sample was divided by the median value for each channel which had itself been divided by the overall dataset median. Differentially abundant proteins were identified using π score, a significance metric that incorporates both fold changes and traditional p-value-based significance scores, determined through a Student’s t-test with or without Welch’s correction, depending on the comparison being made (30).

### Venn, Enrichment, and Network analysis

Venn diagrams were plotted using http://bioinformatics.psb.ugent.be/webtools/Venn/. Enrichment analysis for biological processes, cellular components and molecular functions was performed using Database for Annotation, Visualization and Integrated Discovery (DAVID) v 2021 (31, 32) and Enrichr (33–35). Graphs were constructed using Microsoft Office 365 and GraphPad Prism v 9.0.

### Seahorse XFp Real-Time ATP Rate assay

Seahorse analysis of ATP production was performed using the Seahorse XFp Real-Time ATP Rate assay, following the manufacturer’s instructions. Briefly C1 and E1 cells were seeded in Seahorse XFp Miniplate (Agilent Technologies, Santa Clara, CA, USA) at an optimized concentration of 2×10^4^ cells/well. Infection with *SL* (moi10) was performed for 6 h. At the end of the infection, the media was replaced with the XFp DMEM media supplemented with Glucose (10 mM), Pyruvate (1 mM), and Glutamine (2 mM) and incubated for 1 h in non-CO2 incubator at 37℃. In the meantime, the cartridge was hydrated with Seahorse calibrant for 45/60 minutes in the non-CO2 incubator. Oligomycin (final concentration 1.5 µM /well) and Rotenone/Antimycin (final concentration 0.5 µM/well) (provided by the manufacturers) were loaded into ports A and B of the sensor cartridge and first loaded into the Agilent Seahorse XF HS Mini Analyzer (Agilent Technologies, Santa Clara, CA, USA) for calibration. Following calibration, the cell culture miniplate was loaded into the instrument to measure oxygen consumption rate (OCR), extracellular acidification rate (ECAR), glycolytic ATP, mitochondrial ATP, total ATP, and ATP index according to the manufacturer’s software. The level of glycolytic and mitochondrial ATP after *SL* infection normalized to the level of ATP in the same cell lines without infection. The impact of ELMO1 on ATP production was assessed by comparing the level of ATP in C1 *SL*/C1 UN with E1 *SL*/ E1 UN in both glycolytic and mitochondrial pathways. The Agilent Seahorse Analytics software was used for data analysis.

### Seahorse XF Cell Mito Stress Test for the assessment of mitochondrial respiration

Seahorse analysis of mitochondria respiration was determined through the Seahorse XFp Cell Mito Stress assay, following the workflow provided by the manufacturer’s instructions. C1 and E1 cells were plated and infected with *SL* as described above. The media was replaced with the XFp DMEM medium supplemented with Glucose (10 mM), Pyruvate (1 mM), and Glutamine (2 mM) and incubated for 1 h in non-CO_2_ incubator at 37℃. Following hydration of the cartridge with the Seahorse calibrant, the following inhibitors were injected into the cartridge according to the manufacturers’ instructions: Oligomycin (1.5 µM) in port A, FCCP (1.5 µM) in port B and Rotenone/Antimycin A (0.5 µM) in port C and followed by measurement of OCR, ECAR, basal respiration, maximal respiration, spare respiratory capacity, and ATP production according to the manufacturer’s software. Data analysis was performed with the Agilent Seahorse Analytics software.

### Western Blot

Cells were washed in Phosphate-buffered saline (PBS) and lysed using Radioimmunoprecipitation assay buffer (RIPA) buffer. Mice tissues were homogenized in RIPA buffer and then centrifuged to collect the cell lysates. The lysates were then separated by Sodium dodecyl-sulfate polyacrylamide gel electrophoresis (SDS/PAGE) using a 10% gel, transferred to a polyvinylidene difluoride (PVDF) membrane and were blocked using 5% skim milk/PBS containing 0.1% Tween20 for 1h at room temperature (RT) and then incubated overnight at 4°C with primary antibody for the following targets: BolA Family Member 1/BOLA1 (Cat No 18017-1-AP, Proteintech, Rosemont, IL, USA), ELMO1 (B-7) (Cat No sc-271519, Santa Cruz Biotechnology, Santa Cruz, CA, USA) for C1/E1 cells, and ELMO1 (AB174298-1001, Abcam, Cambridge, United Kingdom) for mouse-derived samples, Dynamin-related protein 1/ DRP1 (12957-1-AP-Proteintech), Sorting Nexin 5/SNX5 (Cat No sc-515215, Santa Cruz Biotechnology, Santa Cruz, CA, USA), Retinoic acid-inducible gene I/RIG-1 (Cat No sc-376845, Santa Cruz Biotechnology, Santa Cruz, CA, USA), Tubulin (Cat No 2144S, Cell Signaling Technology, Danvers, Massachusetts, USA), and GAPDH (Cat No Cell 2118S, Cell Signaling Technology, Danvers, Massachusetts, USA). Anti-mouse IgG, HRP-linked Antibody (Cat No 7076S, Cell Signaling Technology, Danvers, Massachusetts, USA) and anti-rabbit IgG, HRP-linked Antibody (Cat No 7074S, Cell Signaling Technology, Danvers, Massachusetts, USA) were used as secondary antibodies. At least 2-3 independent experiments were performed for each target. Band intensity was assessed using Image J and results were presented as the mean ± SEM. *P values* were determined using unpaired Student’s t-test and were considered significant if the values were < 0.05.

### Functional assay with DRP1-inhibitor Mdivi-1 and ELISA of proinflammatory cytokine

C1 and E1 J774 macrophages (J774) were seeded at a cell density of 5×10^4^ cells/well in a 96-well plate. The following day, cells were infected with *SL* at moi of 10 for 3h and 6h. One hour prior to infection cells were pre-treated with 10 μg/ml of Mdivi1 (Cat # 3982, Bio-Techne Corporation, Minneapolis, MN, USA). For the 6 h time point infection, extracellular bacteria were killed by gentamicin treatment (500 µg/mL) for 90 minutes, followed by treatment with low-dose gentamicin (50 µg/mL) for 3.5 h at 37^◦^C. Mdivi1 was added throughout the entire experiment. Cell supernatants were collected and processed for ELISA. TNF-α was measured using the mouse TNF-α DuoSet ELISA kit (Cat # DY410, R&D, Minneapolis, MN, USA), following the manufacturer’s instructions.

### Infection of mice

Age- and gender-matched WT and global ELMO1 KO C57BL/6 mice were infected with *SL1344* by oral gavage (5×10^7^ cfu/ mouse) as described before (Sayed et al., 2021). The mice were euthanized after 5 days of infection and the ileum were collected for western blot analysis. Animals were bred, housed, used for all the experiments, and euthanized according to the University of California San Diego Institutional Animal Care and Use Committee (IACUC) policies under the animal protocol number S18086.

### Data repository

The dataset files are uploaded to Massive as a private dataset. The link to download them is following (ftp://MSV000091439@massive.ucsd.edu). The DOI for the dataset is following (doi:10.25345/C5M32NM0T), the Massive Identifier number is (MSV000091439) and the Proteome Exchange Identifier number is (PXD040691).

### Statistics

Data reported in this study was analyzed using GraphPad Prism 10 (GraphPad Software Inc., SanDiego, CA, USA). Data is represented as mean ± standard deviation (SD) or standard error of the mean (SEM), otherwise, it is specified. Analysis of data was performed using Student’s t-test or ANOVA one-way test. *p* value < 0.05 was considered significant.

## Results

### The proteomic landscape of macrophages infected with *Salmonella*

To understand the major host pathways linked to ELMO1 after *Salmonella* infection, murine macrophage J774 cells (either control-C1 or ELMO1-depleted E1 cells) were infected with WT *Salmonella* (*SL*), *SifA mutan*t *SL* and compared with the uninfected controls (Fig 1A). The proteomics identified a total of 7777 proteins with 200148 peptide spectral matches in the six conditions with C1 UN, E1 UN, C1 *SL*, E1 *SL*, C1 *SifA*, E1 *SifA* (Fig 1A, Fig S1) with a false discovery rate of 1%. At first, we performed quantitative proteomics and identified the altered proteins in the host cells in uninfected control cells (C1 UN and E1 UN) and cells infected with *SL* (C1 *SL* and EL *SL*) (Fig 1B). The volcano plot showed the deregulated proteins for both statistical significance and fold change. Volcano plots were constructed to identify the most biologically significant proteins (with larger fold change expression) among the pairs mentioned in Fig 1B. We have compared a total of 10 pairs as shown in Table 1. In the pairs P1 ( E1 UN vs C1 UN), P8 (E1 *SL* vs C1 *SL*), P7 (C1*SifA* vs C1 *SL*), P4 (E1 *SifA* vs E1 *SL*), P9 (E1 *SifA* vs. C1 *SifA)* and P10 (E1 *SifA* vs C1 *SL*) 74, 168, 2, 5, 60 and 86 significantly upregulated proteins (blue dots) and 40, 126, 19, 29, 37and 61 significantly downregulated proteins (red dots) were identified, respectively (Table S1-S10). In all E1 cell conditions, ELMO1 protein was lowered compared to C1 cells which provided the confidence of the multiplex proteomics technology and confirmed the characteristic of these ELMO1-depleted macrophages (E1 cells) (Fig 1B).

**Figure 1.**
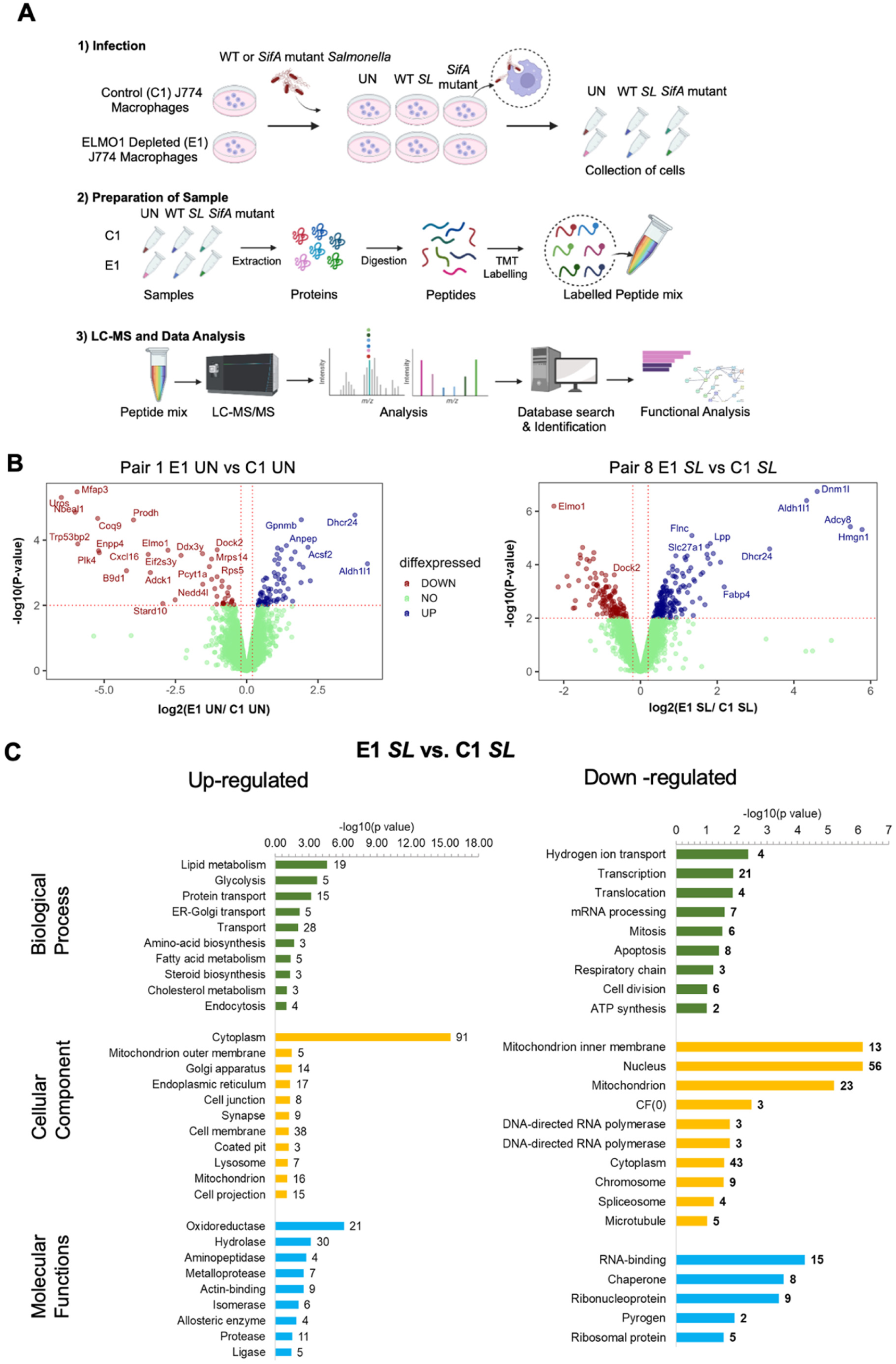
Proteomic profiling and Pathway Enrichment Analysis (PEA) of differentially expressed proteins in control and ELMO1-depleted murine macrophage J774 cells after infection with WT *Salmonella* (*SL*) Diagram showing the main steps in proteome profiling used in this study: (1) infection of control (C1) or ELMO1-depleted shRNA (E1) J774 macrophages with *Salmonella* enterica serovar *typhimurium* strain SL1344 (*SL*); (2) samples preparation, tandem mass tag (TMT) labeling of the pooled sample from the triplicate sets of each condition; (3) separation by liquid chromatography (LC) and mass spectrometry (MS) to quantify relative protein intensities by tags followed by analysis to predict the functions of the differentially expressed proteins. B) Volcano plots showing the statistical significance (log10 *p-value*, Y axis) and fold change of protein expression (log2, X-axis) of different pairs as indicated in the figures. (i) comparison of host proteins of C1 (uninfected) vs host proteins of E1(uninfected). (ii) comparison of host proteins of C1 (*SL*) vs host proteins of E1(*SL*). Red, blue, and green dots represent downregulated, upregulated, and no change in protein levels, respectively. C) GO enrichment and DAVID Functional Annotations for analysis of 293 proteins [upregulated proteins (n=168) and downregulated proteins (n=125)] in C1 *SL* vs E1 *SL*. Green Bars represent biological functions, yellow Bars represent cellular components, and blue bars represent molecular functions. The left panel represent upregulated proteins and the right panel represents downregulated ones.

**Table 1:** List of sample comparisons.

| Pair | Comparison |
| --- | --- |
| 1 | E1 UN vs. C1 UN |
| 2 | E1 SL vs. E1 UN |
| 3 | E1 *SifA* mutant vs. E1 UN |
| 4 | E1 *SifA* vs. E1 SL mutant |
| 5 | C1 SL vs. C1 UN |
| 6 | C1 *SifA* mutant vs. C1 UN |
| 7 | C1 *SifA* vs. C1 SL mutant |
| 8 | E1 SL vs. C1 SL |
| 9 | E1 *SifA* mutant vs. C1 *SifA* mutant |
| 10 | E1 *SifA* vs. C1 SL mutant |

Further, the differentially expressed protein (DEPs) in pair 1 E1 UN vs C1 UN were analyzed by Gene Ontology (GO) enrichment pathway (GO) using the **<u>D</u>**atabase for **<u>A</u>**nnotation, **<u>V</u>**isualization and **<u>I</u>**ntegrated **<u>D</u>**iscovery (DAVID) version 2021. Importantly, the transforming growth factor beta receptor signaling pathway (TGF) and Rac protein signal transduction are among the enriched biological processes in downregulated proteins (Fig S2). It is not surprising, since ELMO1 interacts with DOCK180 and is involved in Rac activation (36). Interestingly, upregulated proteins were identified to be associated with lipid metabolic process, vesicle-mediated and intracellular protein transport, and filament organization and assembly (Fig 1C, Fig S2). These finding allign with previous findings showing that ELMO1 plays a role in the filament assembly and endocytosis/ engulfment of bacteria (18, 37, 38). After infection with *SL*, functional annotation of significantly upregulated proteins from E1 *SL* vs C1 *SL* identified also major proteins linked with lipid metabolism, transport, apoptosis, and ATP synthesis (Fig 1C) while transcription, apoptosis, and hydrogen ion transport were some of the major functions associated with downregulated proteins (Fig 1C). Importantly, the protein-protein interaction (PPI) networks for DEPs showed many pathways linked to mitochondrial activity such as oxidative phosphorylation, metabolic pathways, integration of energy metabolism, chemokine signaling pathways, and Inositol phosphate metabolism (Fig S3).

### Alteration of mitochondrial functions in ELMO1-depleted macrophages compared to control during *Salmonella* infection

To identify the proteins involved with ELMO1 during *SL* infection, we plotted a Venn diagram of upregulated and downregulated proteins from pairs 1, 2, 5, and 8 where C1 and E1 cells either uninfected or infected with *SL* were compared together (Fig 2A). Comparison of DEPs from pairs 1, 2, 5, and 8 identified 153 (pair 2), 91 (pair 8), and 19 (common to pair 2 and 8) upregulated unique proteins in E1 cells after *SL* infection. Besides, there were 44 (pair 2), 105 (pair 8), and 11 (common to pair 2 and pair 8) proteins were downregulated in E1 cells during *SL* infection (Fig 2B). Although 153 (pair 2 upregulated) and 44 (pair 2-downregulated) proteins were identified as DEPs these proteins were compared to E1 uninfected. Since we were interested in DEPs in E1 cells compared to C1 cells during *SL* infection, we further evaluated 19 upregulated proteins and 11 downregulated proteins that are common in both pair 2 and pair 8 (Fig 2B). GO enrichment analysis using Enrichr revealed that the 19 upregulated and the 11 downregulated proteins are involved in various functions predominantly linked with immune response, cell signaling, and mitochondrial pathway. Other cellular functions associated with the upregulated proteins include vesicle-mediated transport, metabolism, and chromatic organization (Fig 2C upper right panel, Table 2) while protein synthesis, DNA replication, and transcription were enriched among the downregulated proteins (Fig 2C lower right panel, Table 3).

**Figure 2.**
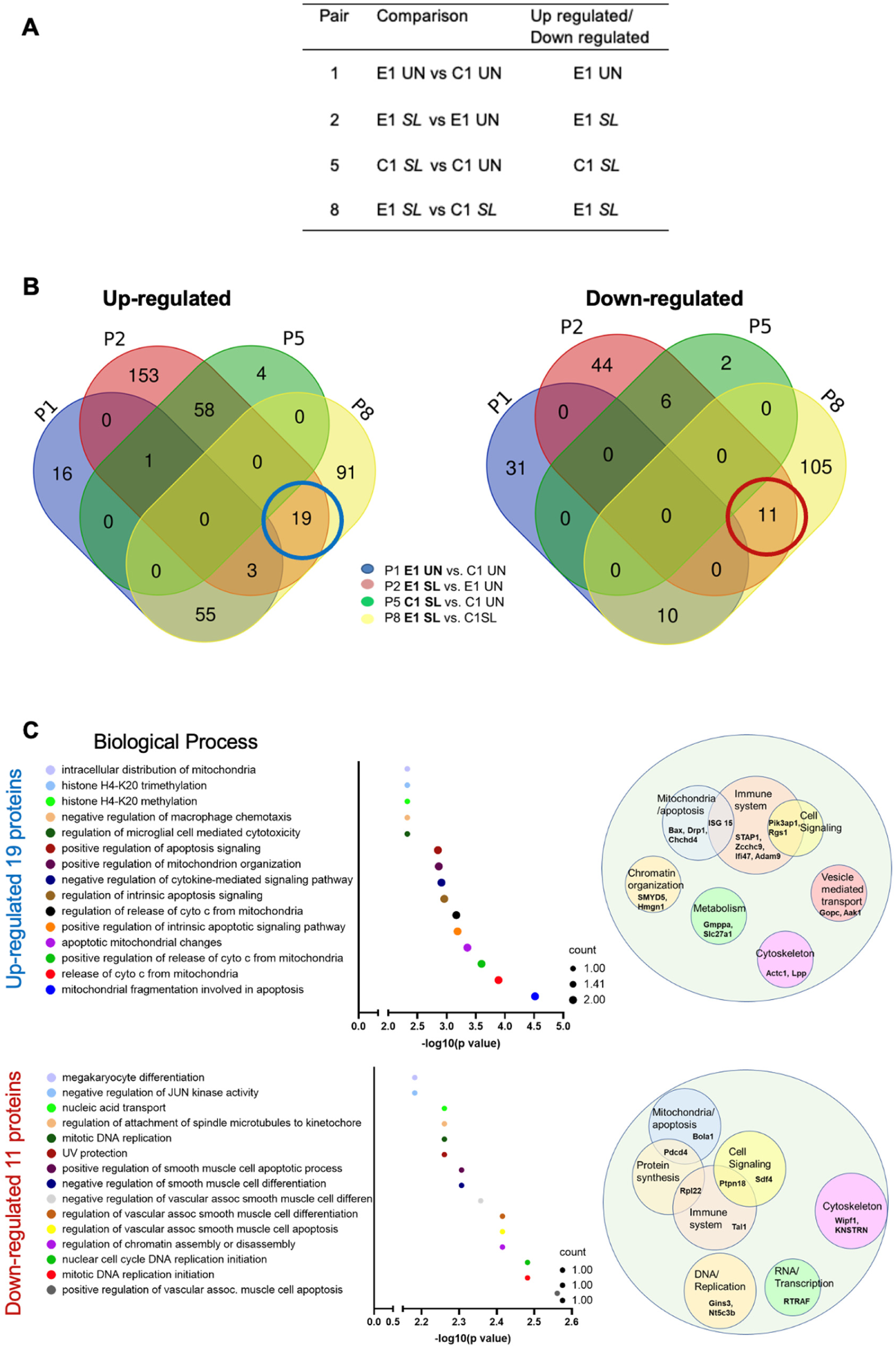
ELMO1 regulates the pathways involved in mitochondrial function, vesicular transport, and host immune responses following WT *Salmonella* infection. A) Table showing the pairs and the respective proteomic datasets considered for the comparisons-Pair 1 (E1 un versus C1 un), pair 2 (E1 *SL* versus E1 un), pair 5 (C1 *SL* versus C1 un), and pair 8 (E1 SL versus C1 *SL)*. B) Venn Diagrams showing the comparison of significantly upregulated proteins (Left) and downregulated proteins (Right) from the above-mentioned pairs. C) (left panel) GO term enrichment analysis showing the biological processes of the 19 upregulated [circled “blue” in Venn diagram (B)] and 11 downregulated proteins [circled “red” in Venn diagram (B)] unique in E1-*SL* conditions (between P2 and P8 where E1 SL is compared with E1 un and C1 *SL*). The size of the circle in the graph corresponds to the number of proteins counts in each pathway. (Right panel) represents the major functions associated with these proteins by using PubMed and uniport search (last accessed on Dec 15, 2023). The size of the circles is arbitrary depending on the number of proteins associated with the same function.

**Table 2.**
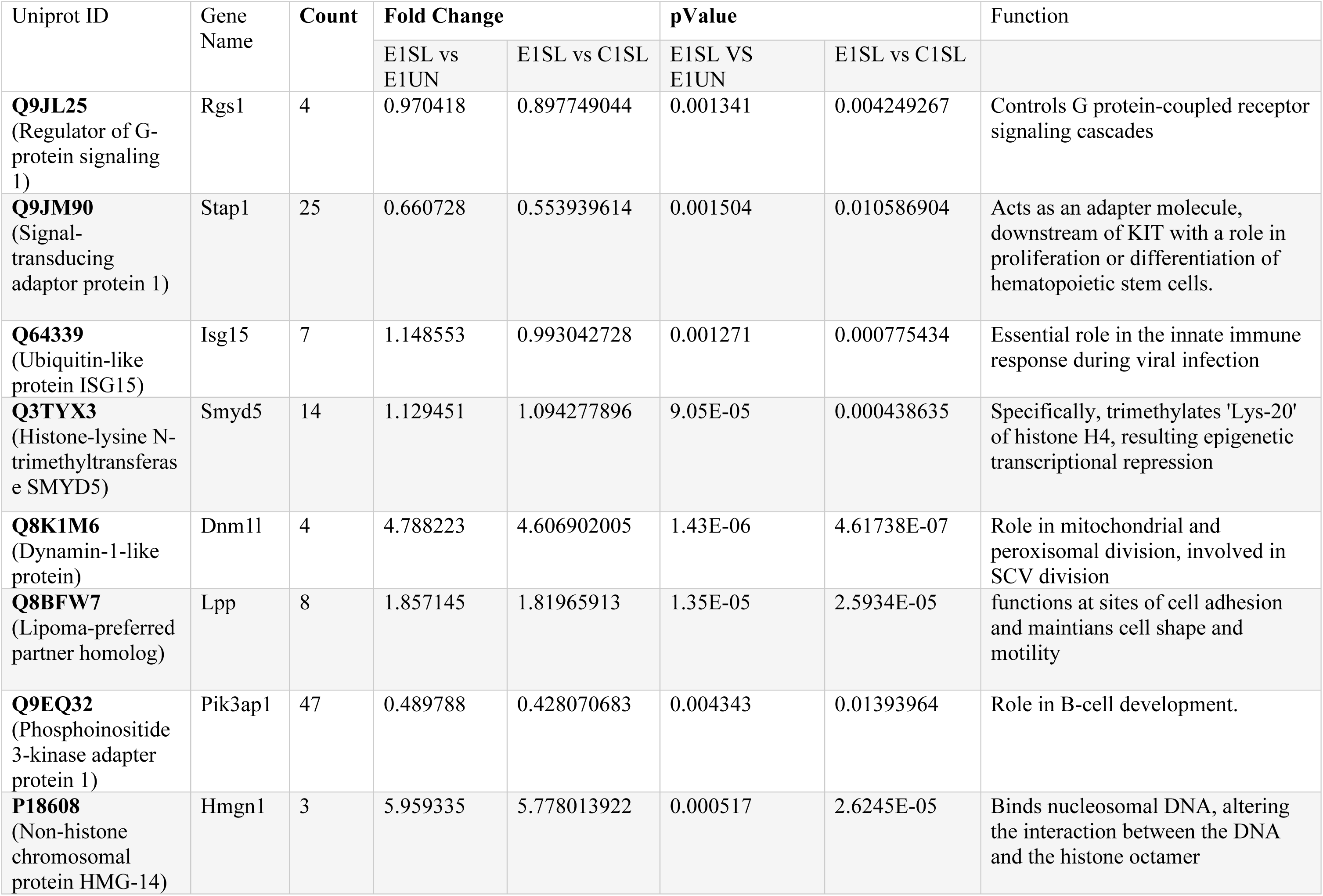
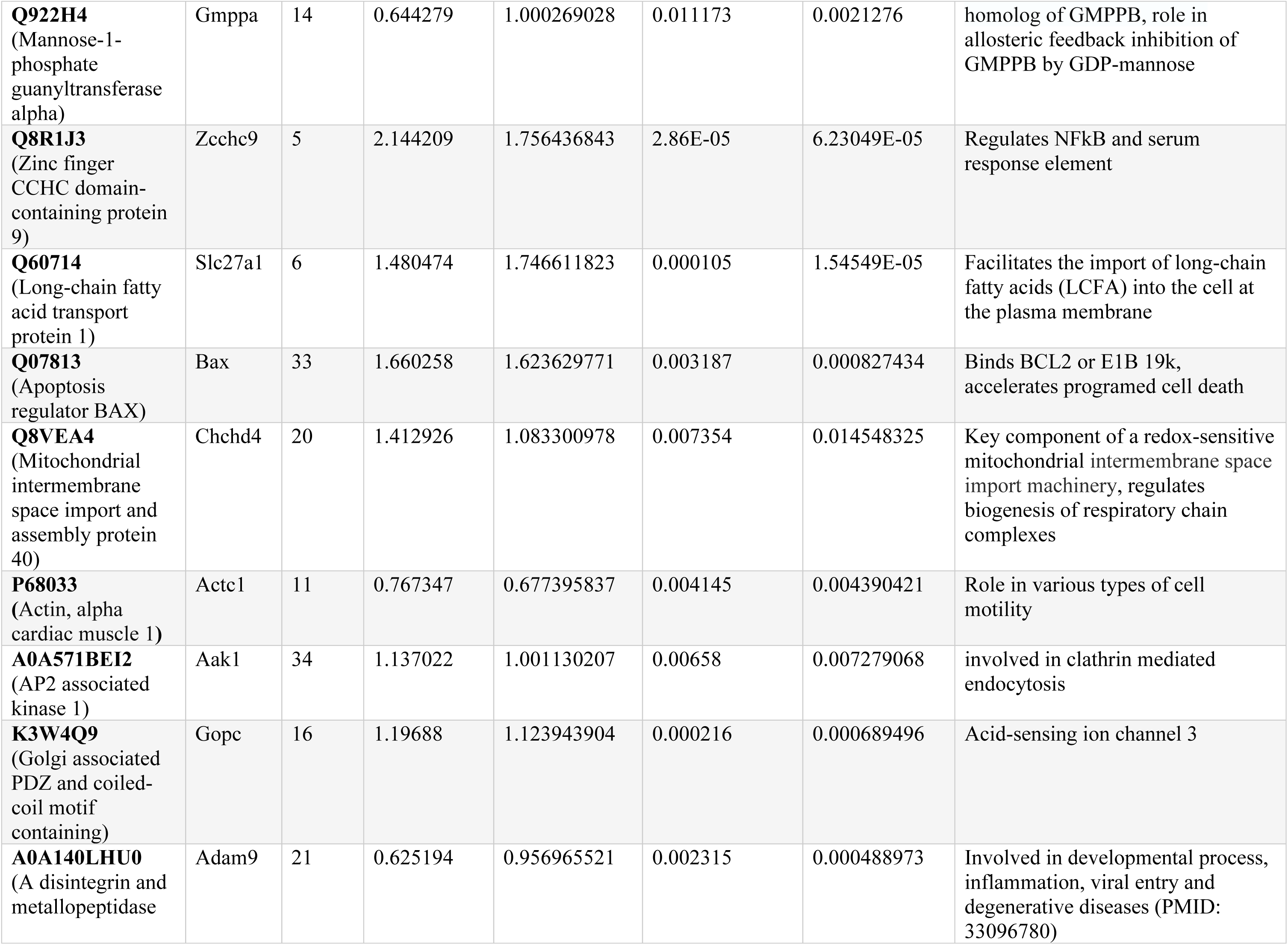
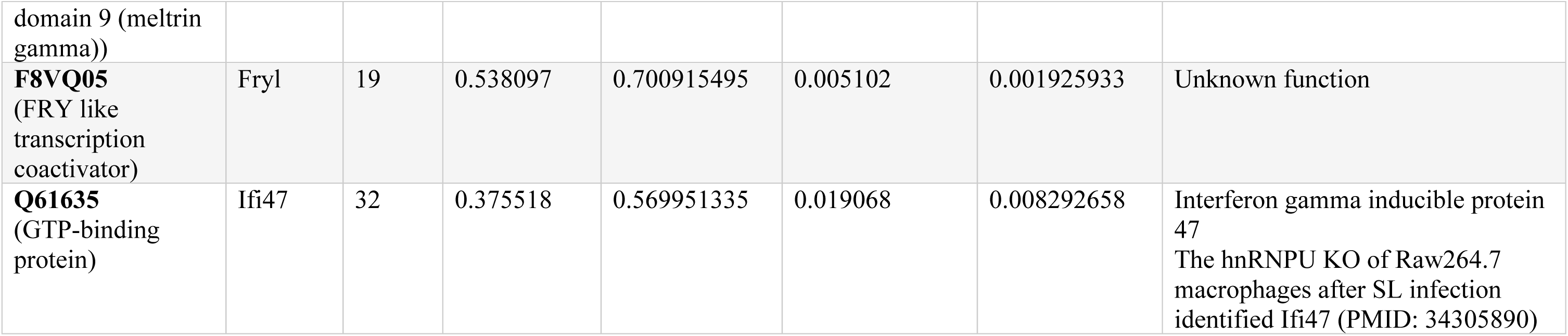
Functions of 19 upregulated proteins identified in E1-SL conditions.

**Table 3.**
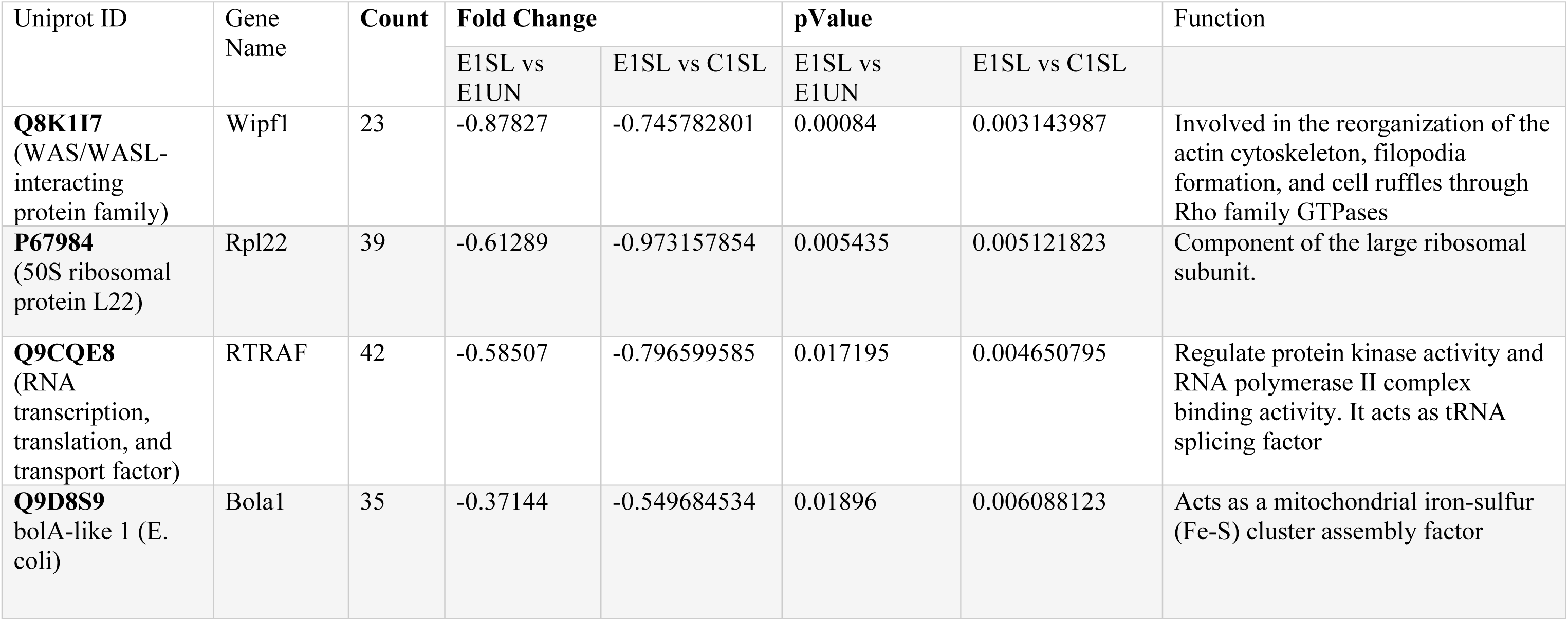
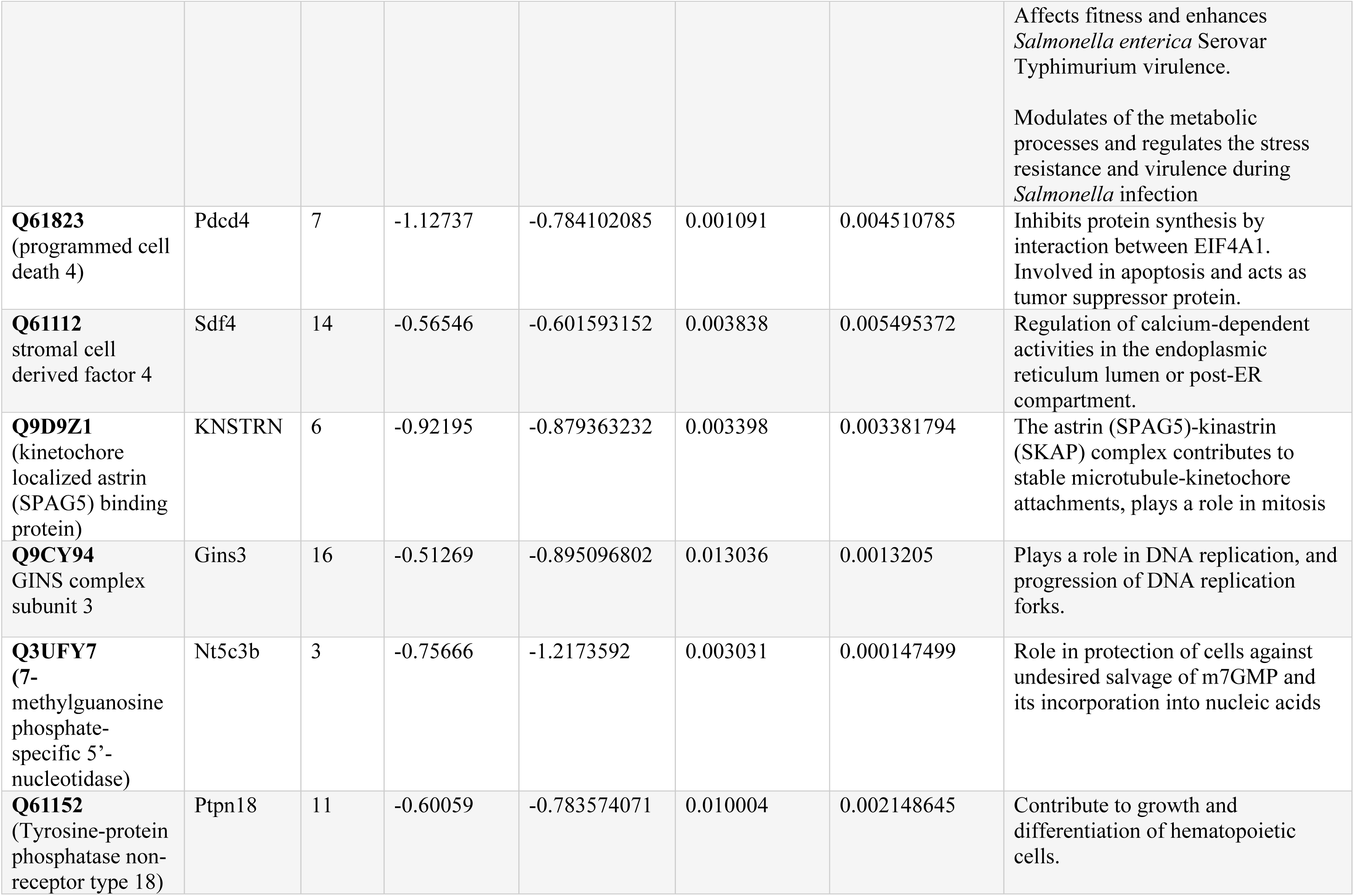
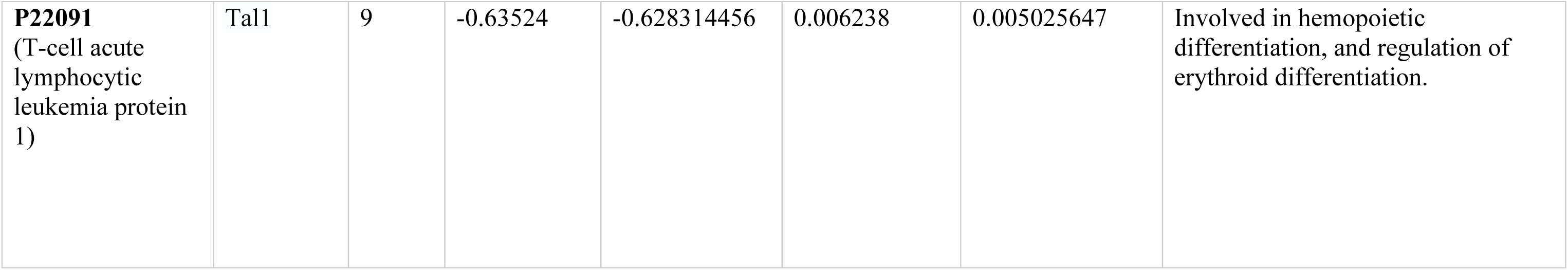
Functions of 11 downregulated proteins identified in E1-SL conditions.

### ELMO1 affects glycolysis and oxidative phosphorylation following *Salmonella* infection

To investigate whether ELMO1 affects the host metabolism during *Salmonella* infection, C1 and E1 cells were infected with *SL* for 6 h, followed by measurement of oxygen consumption rate (OCR) and the extracellular acidification rate (ECAR), as depicted in Fig 3 A-C, using the Seahorse XFp real-time ATP rate assay. We found that *SL* infection induces a significant increase in glycolysis and a significant decrease in oxidative phosphorylation (OXPHOS) (Fig 3 B-C, Fig S4), indicating a metabolic shift in macrophages from OXPHOS to glycolysis upon *SL* infection, a phenomenon previously observed in other studies (39–44). Importantly, we observed that the changes induced by *SL*, are exacerbated by the depletion of ELMO1 as reflected by the increased percentage of glycolysis (Fig 3D) and a further reduction in the percentage of oxidative phosphorylation (Fig 3E) in E1 cells when compared to C1 macrophage during *SL* infection. Moreover, our results indicate a reduction of the ATP rate index, which represents the ratio of the mitochondrial ATP to the glycolytic ATP production, in E1 cells following *SL* infection (Fig 3F, Fig S4). Next, we performed the Seahorse XFp Cell Mito Stress assay to determine the impact of ELMO1 on aerobic respiration. We confirmed a reduction in OCR, indicating an overall decrease in mitochondrial respiration (Fig 3G) during infection. Calculated values derived from the Seahorse XFp analyzer demonstrated a significant decrease in mitochondria ATP production (Fig 3H), basal respiration (Fig 3I), maximal respiration (Fig 3J), and spare respiratory capacity (Fig 3K) in C1 and E1 cells following *SL* infection.

**Figure 3.**
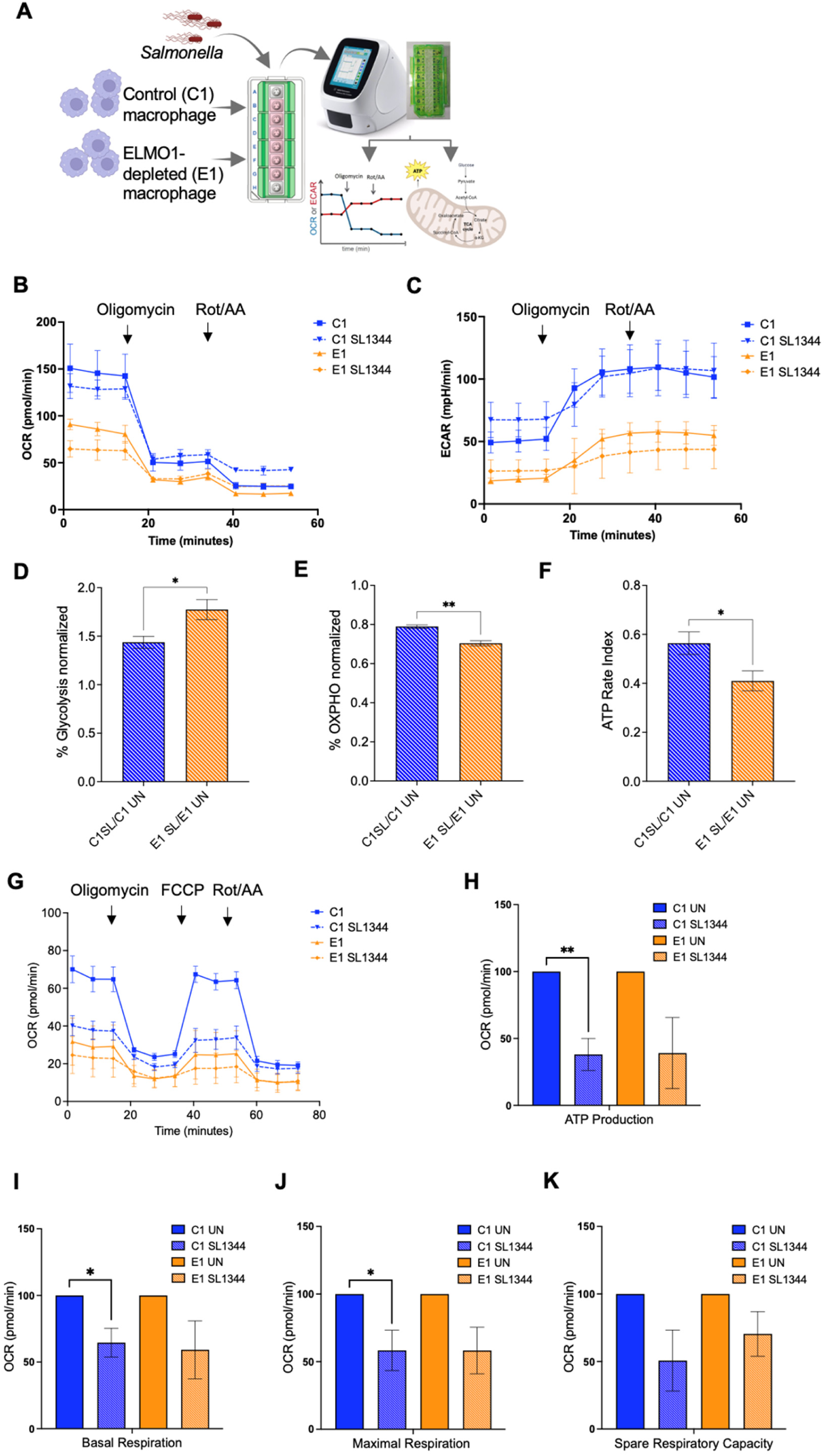
The functional role of ELMO1 in mitochondrial dynamics following infection with WT *Salmonella* by bioenergetic analysis and ATP production. A. Schematic representation of the experimental workflow for the Seahorse assays. Control shRNA J774 cells (C1) and ELMO1-shRNA J774 (E1) were seeded in XF HS Miniplate and challenged with WT SL1344 strain (*SL*) or left untreated. After 6 h, the plate was prepared for the ATP rate or Mito Stress assay, and the assay was run using the Agilent Seahorse XF HS Mini Analyzer. B. Representative kinetic profile of Oxygen Consumption Rate (OCR) measurements (pmol/min) presented as averages of three individual replicates per condition ± standard deviations in C1, E1 at basal level or following 6 h infection with *Salmonella* (*SL*). Data presented as averages of three technical replicates. C. Representative kinetic profile of Extracellular Acidification Rate (ECAR) measurements (mpH/min) in C1, E1 at basal level or following 6 h infection with *SL*. The assay was repeated three times and was performed in triplicates using the Seahorse Xfp real-time ATP rate assay. Data presented as averages of three technical replicates. D-E. Quantification of percentage of glycolysis (D) and. Oxidative Phosphorylation (E) in C1 and E1 following *SL* infection for 6h as shown in B and C. Values were normalized to the respective values in uninfected C1 and E1. F. Quantification of ATP rate index (Ratio of the mitochondria ATP divided by glycolytic ATP Production) in C1 and E1 following *SL* infection for 6 h as in B, C, and D. Values were normalized to the respective C1 E1 untreated values. G. Representative kinetic profile of Oxygen Consumption Rate (OCR) (pmol/min) of Seahorse XF Cell Mito Stress Test assay in C1 and E1 cells at basal level or following 6 h infection with SL. Data presented as averages of three technical replicates. H-K. The OCR of (H) ATP production, (I) basal respiration, (J) maximal respiration, and (K) spare respiratory capacity of experiments in G were determined. Data presented as averages of three independent experiments, where * indicates p value ≤ 0.05 and ** indicates p value ≤ 0.01 as assayed by two-tailed Student’s t-test.

### Validation of DEPs in ELMO1-depleted macrophages after *Salmonella* infection

To validate the proteomics data and the effect of ELMO1 on the identified proteins, we performed *in vitro* and *in vivo* infection experiments and assessed the expression of selected proteins by western blot. We selected one downregulated protein; BolA Family Member 1 (BOLA1) and one upregulated protein, dynamin-related protein 1 (DRP1) from the selected list based on their involvement in mitochondrial function. Among the 11 downregulated proteins, we selected BOLA1 as a mitochondrial protein that affects mitochondrial morphology, protects the cells from oxidative stress, and influences other mitochondrial processes (45, 46). *In vitro* experiments data showed that the expression levels of BOLA1 were decreased after *SL* infection, and it was significantly lower in E1 *SL* compared to all uninfected cells (Fig 4A).

**Fig 4:**
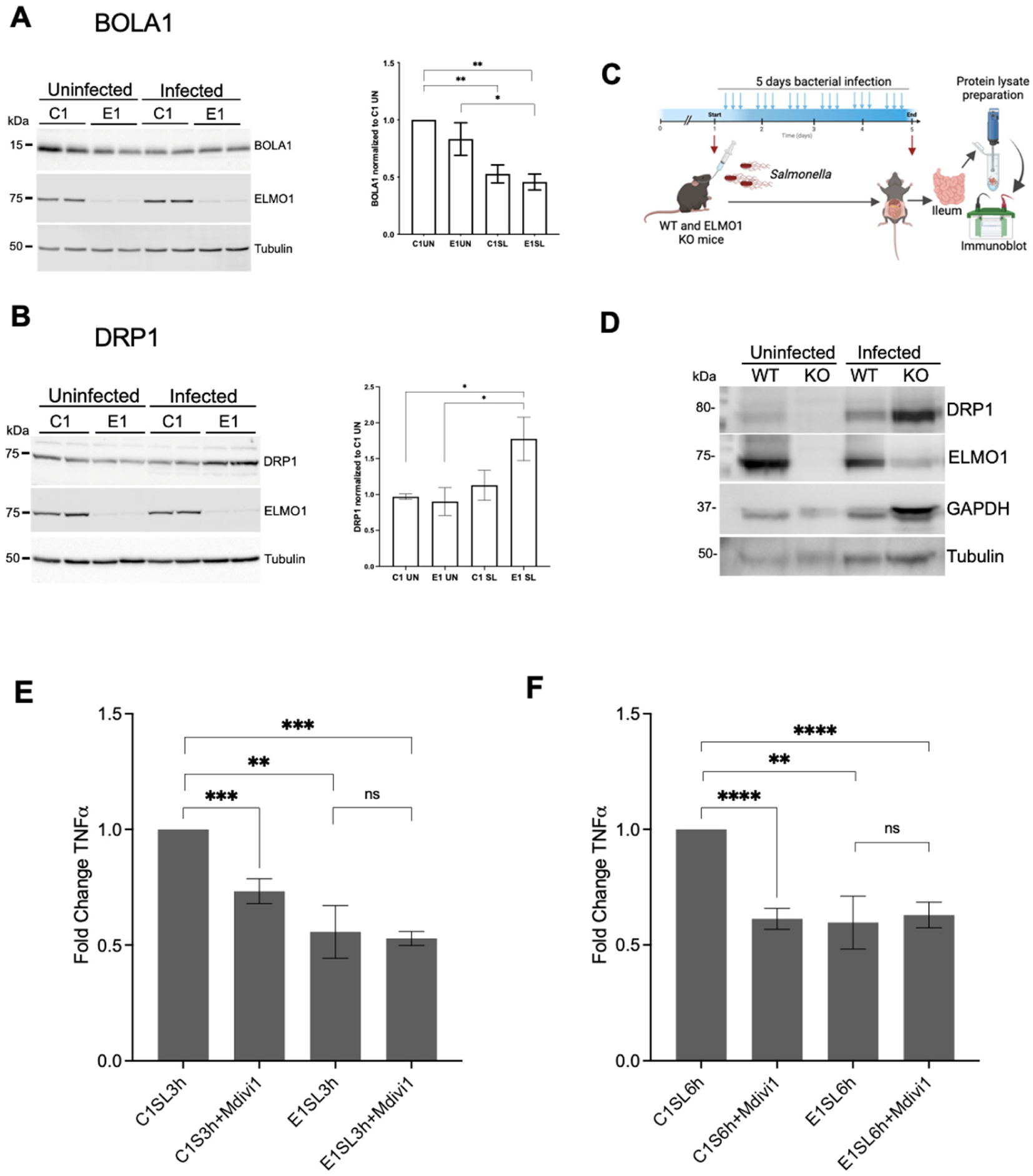
Validation of targets by western blots and functional analysis. A-B Western Blot of BOLA1 (A) and DRP1 (B) in C1-E1 cells following infection with *SL*. The densitometry was performed using three independent experiments where * indicates p value ≤ 0.05 and ** indicates p value ≤ 0.01 as assayed by two-tailed Student’s t-test. C. Schematic illustration on animal experiments for the assessment of the expression of DRP1 as in D. D. Western blot of DRP1 using the tissue samples from the Ileum of WT and ELMO1 KO mice after 5 days of *SL* infection, as represented in C. E-F. TNF-α cytokine levels in the supernatant from C1 and E1 cells after 3h (E) and 6h (F) infection with *SL* in the presence of DRP1 inhibitor mDivi-1, measured by ELISA. Data represents fold change (mean ± SEM) compared to C1 from the values collected from two separate experiments with four biological replicates. * Indicates p ≤ 0.05, ** indicates p ≤ 0.01, *** indicates p ≤ 0.001, **** indicates p ≤ 0.0001 as assayed by two-tailed Student’s *t*-test.

On the other hand, DRP1 which is a member of the dynamin superfamily of GTPases and involved in mitochondrial fission (47, 48) was found to be upregulated in E1 *SL* compared to other conditions. We noticed that the expression of DRP1 was significantly upregulated in E1 *SL* compared to all uninfected conditions as well as when compared to C1 *SL* (Fig 4B). To further confirm our findings, we assessed the level of DRP1 in the ileum of wild-type and ELMO1-KO mice infected with *SL* for 5 days (Fig 4C). Interestingly, we found that the highest level of DRP1 was detected in the ileum of *SL*-infected ELMO1-KO mice (Fig 4D).

### Functional analysis of DRP1 in inflammation

Previous studies have shown that DRP1 modulates the production of inflammatory cytokines in response to immune stimulation and infection (49, 50). To understand whether the role of ELMO1 is linked to the function of DRP1, we have used the DRP1 inhibitor Mdivi-1(51) in C1 and E1 cells following infection with *SL* for 3h (Fig 4E) and 6h (Fig 4F). The level of TNF-α is downregulated in E1 compared to C1 before Mdivi-1 treatment. TNF-α levels were significantly reduced in *SL*-infected C1 cells following Mdivi-1 treatment, while it did not decrease further in E1 cells after infection (Fig 4E, Fig 4F and Supplement figure S5). This finding suggests that the majority of the DRP1-dependend TNF-α response is mediated by ELMO1.

### Pathway enrichment analysis of DEPs controlled by ELMO1 after infection with *SifA* mutant

Previously, we have shown that ELMO1 interacts with the *Salmonella* effector SifA (24). Here, we determined the DEPs after *Salmonella* effector protein SifA during infection in the presence and absence of ELMO1. We performed the analysis of the proteins in pair 7 ( C1 *SifA* vs C1 *SL*), pair 4 (E1 *SifA* vs E1 *SL*), pair 9 (E1 *SifA* vs C1 *SifA*), and pair 10 (E1 *SifA* vs C1 *SL*). A comparison of proteins from pair 7 revealed that there are 19 downregulated proteins and 2 upregulated proteins myosin light chain peptide 6 (Myl6) and Plasminogen activator urokinase receptor PLAUR (Fig 5A). The two upregulated proteins are involved in cell proliferation, migration, and apoptosis (PLAUR) or motor protein myosin (My16). Similarly, a comparison of pair 4 proteins identified 30 and 5 proteins with elevated expression levels in the case of *SL* and SifA mutant infection (Fig 5B). The 5 upregulated proteins are involved in the cellular processes (Rpl11, NudE, and Mat2a) and linked to Myc pathways (Ndrg2 and Mycbp).

**Fig 5.**
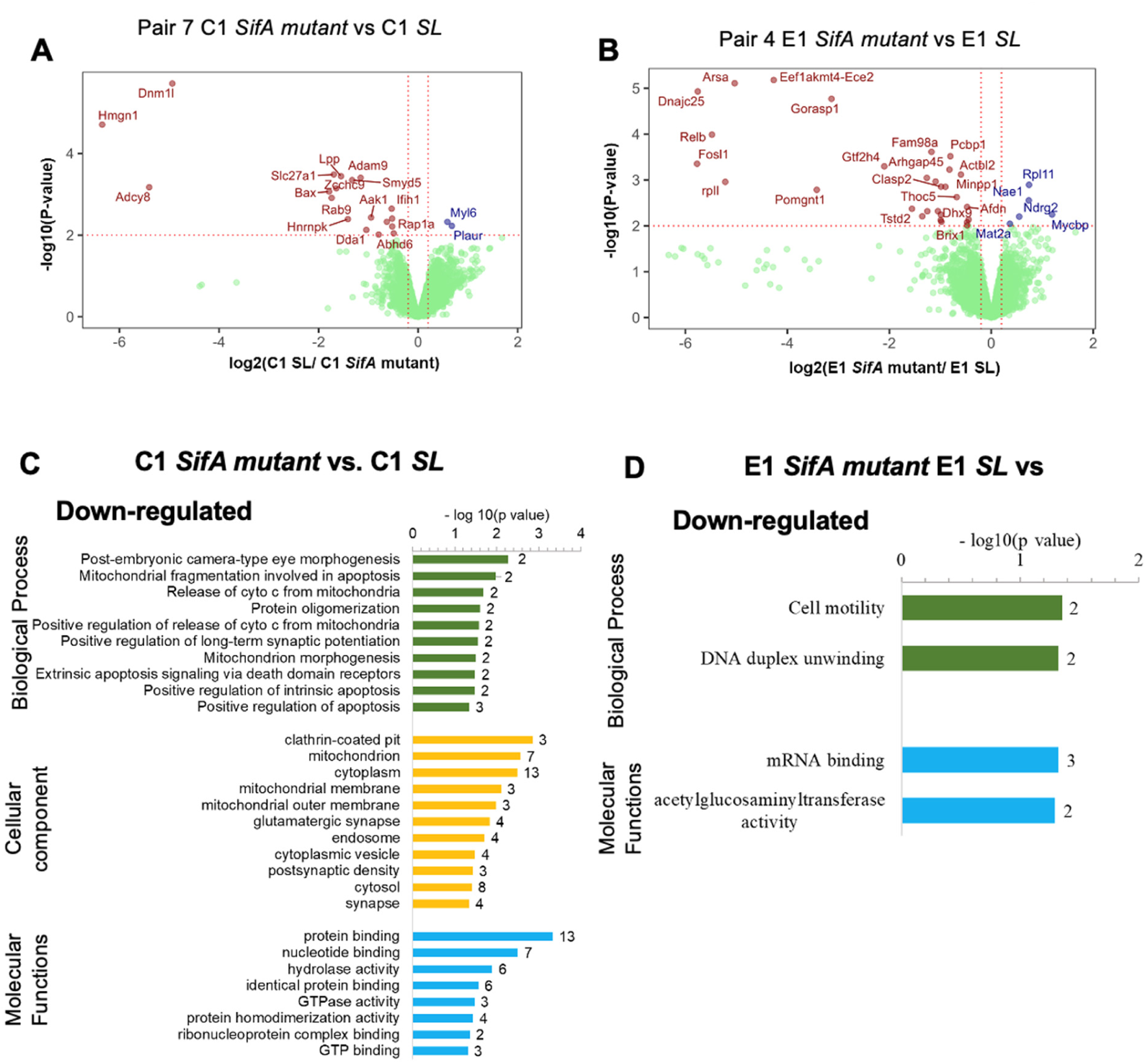
PEA and DAVID analysis for control shRNA (C1) and ELMO1 shRNA (E1) in J744 cells after infection with the *SifA* mutant strain. A & C. Volcano plots showing the statistical significance (log10 *p-value*, Y axis) and fold change of protein expression (log2, X-axis) of different pairs as indicated in the figures. Pair 7 C1 *SifA* mutant vs C1 *SL* in A and Pair 4 E1 *SifA* mutant vs E1 *SL* in C. Red, blue, and green dots represent downregulated, upregulated, and no change in protein levels, respectively. B & D. GO term enrichment analysis and DAVID functional annotations for DEPs of biological functions (green), cellular component (yellow), and molecular functions (blue) of Downregulated proteins of Pair 7 (C1 *SifA* mutant vs C1 *SL*) in B and Downregulated proteins of Pair 4 (E1 *SifA* mutant vs E1 *SL*) in D.

The GO analysis of the downregulated proteins in C1 *SifA* vs C1 *SL* from pair 7 (Fig 5A) highlighted pathways involved in mitochondrial functions and apoptosis, protein oligomerization, and embryonic development functions (Fig 5C). GO analyses of proteins in E1 *SifA* vs E1 *SL* in pair 4 (Fig 5B) revealed that cell motility and DNA duplex unwinding are the significant functions associated with downregulated E1 proteins during *SifA* infection (Fig 5D).

Next, we checked the effect of ELMO1 and SifA, in pair 9 *SifA mutant* infection (Fig 6A) of C1 and E1 cells and in pair 10 with E1 *SifA* vs C1 *SL* (Fig 6B). The proteins in Fig 6A are linked with metabolic functions such as glucose, lipid, and cholesterol were higher in E1 cells compared to C1 whereas proteins involved in immune response, cytokine response mediated by NFĸB and stress response had lower expression levels (Fig 6C). Similarly, a comparison of pair 10 with E1 *SifA* vs C1 *SL* (Fig 6B) could detect 86 proteins with higher levels in E1 after SifA mutant infection. Functions associated with metabolism (lipid, glycolysis, and sterol) and ER-Golgi transport were the most enriched biological processes (Fig 6D). Also, 61 proteins had higher expression levels in C1 cells infected with *SL,* and cell cycle and mitosis were the prominent biological processes identified upon enrichment analysis (Fig 6D). Among various pathways and functions that are controlled by ELMO1 during *SL* and *SifA mutan*t infection, we noticed that most of these pathways are associated with metabolism and host immune/defence responses.

**Figure 6.**
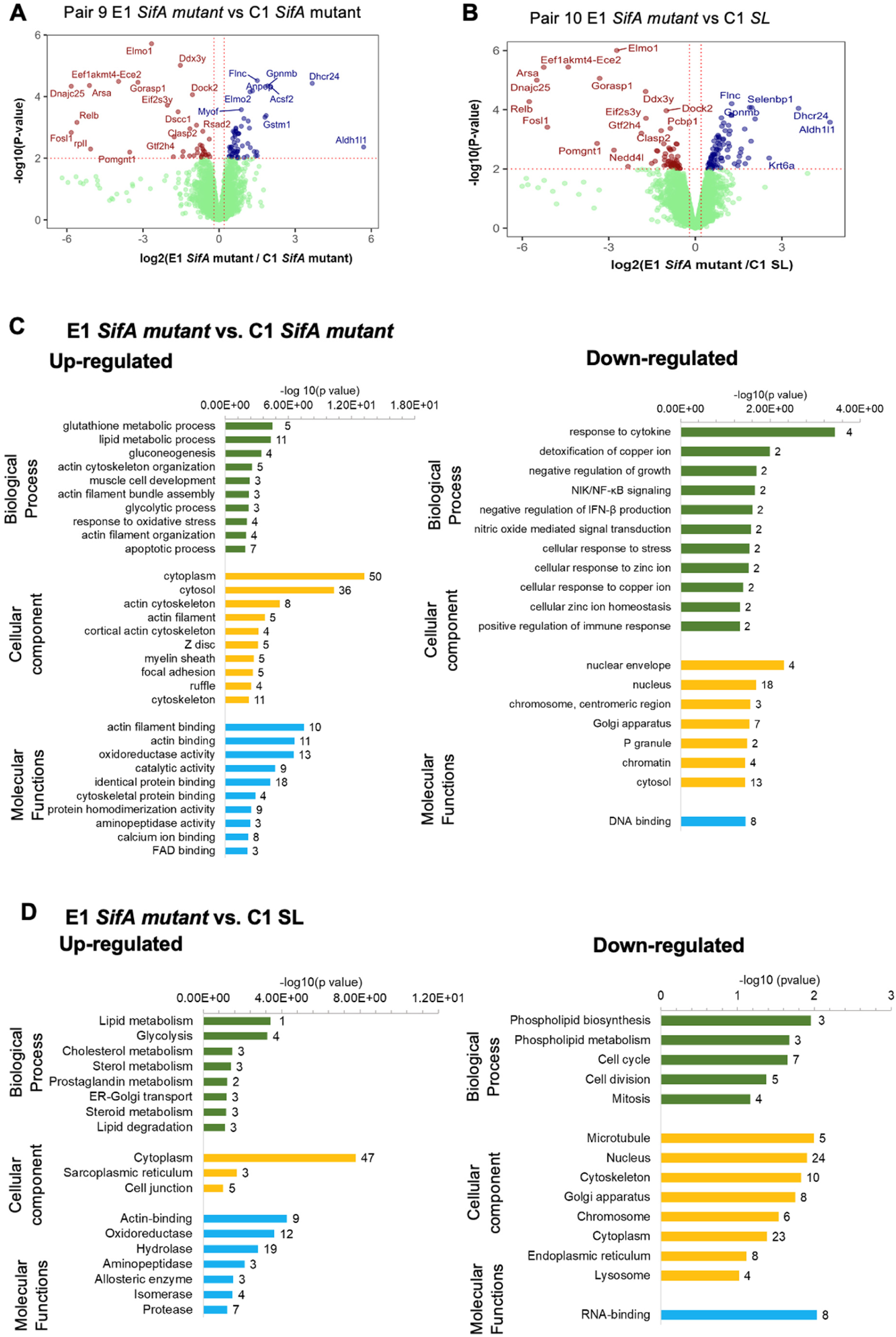
PEA and DAVID analysis for control shRNA (C1) and ELMO1 shRNA (E1) J744 cells after infection with WT *SL* and *SifA* mutant strain. A & C. Volcano plots show the statistical significance (log10 *p-value*, Y axis) and fold change of protein expression (log2, X-axis) of different pairs as indicated in the figures. Pair 9 E1 *SifA* mutant vs C1 *SifA* mutant in A and Pair 10 E1 *SifA* mutant vs C1 *SL* in C. B & D. GO term enrichment analysis and DAVID functional annotations for DEPs of biological functions (green), cellular component (yellow), and molecular functions (blue) of: (B) Upregulated proteins (left) and downregulated proteins (right) of Pair 9 E1 *SifA* mutant vs C1 *SifA* mutant) and (D) Upregulated proteins (left) and downregulated proteins (right) of Pair 10 (C1 *SL* vs E1 *SifA* mutant).

### The differentially regulated proteins controlled by ELMO1-SifA interaction

To identify the proteins involved in *SifA* infection in presence and absence of ELMO1, we plotted Venn diagram of upregulated and downregulated proteins from the following pairs: pair 1 (E1 Un vs C1 un), pair 7 (C1 *SifA* vs C1 *SL*), pair 8 (E1 *SL* vs C1 *SL*), pair 9 (*E1 SifA vs* C1 *SifA)*, and pair10 (*E1 SifA* vs C1 *SL)* (Fig 7A). Accordingly, 11 proteins with higher expression levels and 18 proteins with lower expression levels were found to be unique in E1 infected with *SifA* mutant (Fig 7B).

**Figure 7.**
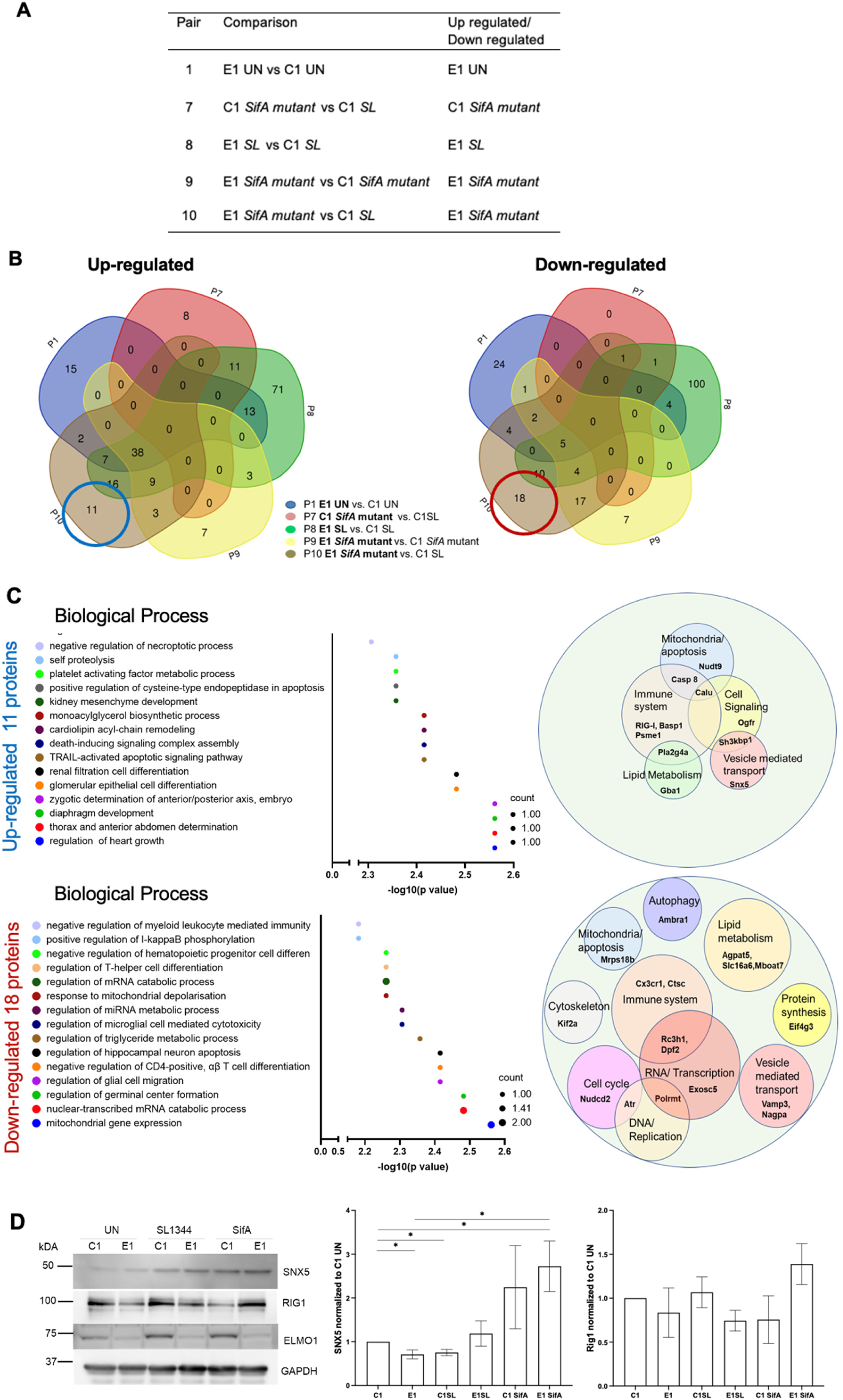
The differentially expressed protein profiles are regulated by host ELMO1 and *Salmonella* effector SifA. A. Table showing the pairs and the respective proteomic datasets considered for the comparison: Pair 1 (C1 Un vs E1 Un), Pair 7 (C1 *SL* vs C1 *SifA* mutant), Pair 8 (C1 *SL* vs E1 *SL*), Pair 9 (C1 *SifA* mutant vs E1 *SifA* mutant), and Pair 10 (C1 *SL* vs E1 *SifA* mutant). B. Venn Diagram comparing significantly upregulated proteins (Left) and downregulated proteins (Right) from the above-mentioned pairs in (A). C. (left) GO term enrichment analysis for biological processes of DEPs in E1 *SifA* compared to other conditions in the above-mentioned pairs. 11 upregulated protein (Upper panel circled blue in Venn diagram B) and 18 downregulated (lower panel circled red in Venn diagram B). The size of the circle in the graph corresponds to the number of protein counts in each pathway. (Right) represents the major functions associated with these proteins by using PubMed and uniport search (last accessed on Jan 15, 2023). The size of the circles is arbitrary depending on the number of proteins associated with the same function. D) Western blot analysis of vesicle transport protein SNX5 and immune system protein RIG1 in C1-E1 cells following infection with *SL*. The densitometry was performed using three independent experiments where * indicates p<0.5 as determined by two-tailed Student’s t-test.

Immune response, mitochondrial functions/apoptosis, lipid metabolism, and vesicle-medited transport are the most enriched GO Terms associated with both the upregulated and the downregulated proteins (Fig 7C, Table 4, and Table 5). Other enriched pathways associated with the downregulated targets include autophagy, DNA replication, and transcription (Fig 7C and Table 5). As immune response and vesicle-mediated transport emerged as common enriched pathways in both upregulated and downregulated proteins, we chose to validate the proteomics findings by focusing on two targets from these pathways: SNX5 and RIG1. SNX5, known as sorting nexin 5, plays a role in endosomal trafficking via interaction with the retromer complex (VPS35, VPS26, and VPS29) (52). RIG-I (retinoic acid-inducible gene-I), is a cytosolic viral RNA sensor with a role in the activation of innate immunity (53). The protein level of SNX5 was increased in E1 cells after *SL* and *SifA* infection, and the highest level was detected in *SifA* mutant infected E1 cells (Fig 7D). Similarly, *SifA* mutant-infected E1 cells showed the highest expression of RIG-1 (Fig 7D).

**Table 4.** Functions of 11 upregulated unique proteins to E1-*SifA* conditions.

| Uniprot ID/ Protein | Gene | Count | Fold Change | pValue | Function |
| --- | --- | --- | --- | --- | --- |
| **Q6XLQ8**<br>(Calumenin) | Calu | 52 | 0.861832837 | 0.01268679 | Contributes to ER-Ca<sup>2+</sup> homeostasis<br>Calumenin is developed during proteomics analysis of *Salmonella* Typhimurium infection |
| **Q6Q899**<br>(Orthologous to human DDX58 (DEXD/H-box helicase 58)) | Rigi<br>RNA<br>sensor<br>RIG-I | 92 | 0.624993687 | 0.035331374 | RIG-I, together with RNA polymerase III, sense viral DNA.<br>RIG-I Detects mRNA of Intracellular *Salmonella* Typhimurium in nonimmune cells |
| **P47713**<br>(Cytosolic phospholipase A2.(PLA2)) | Pla2g4a | 48 | 0.469264737 | 0.038163132 | Plays a role in allergic reactions, parturition, fertility, neuronal death,<br>membrane lipid remodeling and production of inflammatory mediators. |
| **Q8BVU5**<br>(ADP-ribose pyrophosphatase, mitochondrial) | Nudt9 | 31 | 0.560039044 | 0.024352214 | Hydrolysis of ADP-ribose to AMP and ribose 5'-phosphate. |
| **Q6P7W2**<br>(SH3 domain-containing kinase-binding) | Sh3kbp1 | 3 | -0.104954062 | 0.81920896 | Prevents CBL-SH3KBP1 complex facilitated down-regulation of EGFR signaling. Adapter protein involved in endocytosis and lysosomal degradation signaling pathways. |
| **A0A087WQT6**<br>Caspase-8 | Casp8 | 41 | 0.586637451 | 0.032543796 | Thiol protease, acts as a molecular switch for apoptosis, necroptosis and pyroptosis; *Salmonella* infection recruit's caspase-8 to the inflammasome and regulate interleukin-1β production; *Salmonella* Typhimurium sseK3 induces caspase 8 activation and modulates macrophage apoptosis and glycolysis. |
| **Q9D8U8**<br>(Sorting nexin-5) | Snx5 | 90 | 0.537662356 | 0.032847912 | Plays a role in various stages of intracellular trafficking; Essential for macropinocytosis and antigen processing; Activates endosomes and autophagy. SNX5 has been identified during proteomic analysis of RAW 264.7 macrophages infected with *Salmonella*. |
| **Q91XV3**<br>(Brain acid soluble protein 1) | Basp1 | 74 | 0.824853969 | 0.009652265 | Interferes with the oncogenic capacity of MYC and its binding to calmodulin<br>Proteomics analysis of Hela cells infection with *Salmonella* revealed that Basp1 is one of *Salmonella* ubiquitin ligase SlrP on host cells. |
| **Q99PG2**<br>(Opioid growth factor receptor) | Ogfr | 46 | 0.636973632 | 0.024963872 | Alternatively known as Met-enkephalin, has a role in growth regulation |
| **P17439**<br>(Lysosomal acid glucosylceramidase) | Gba1 | 24 | 0.581964936 | 0.020146627 | Increase cholesterol glucosylation activity. Mutation of Gba1 is associated with Parkinson disease; catalyzes the hydrolysis of glucosylceramides within the lysosomal compartment |
| **P97371**<br>(Proteasome activator complex subunit 1) | Psme1 | 78 | 0.492696198 | 0.036703895 | PA28 modulates antigen processing of viruses; Role in immunoproteasome assembly and essential for efficient antigen processing; *Salmonella* Typhimurium increased the expression of PA28 proteasome genes |

**Table 5.**
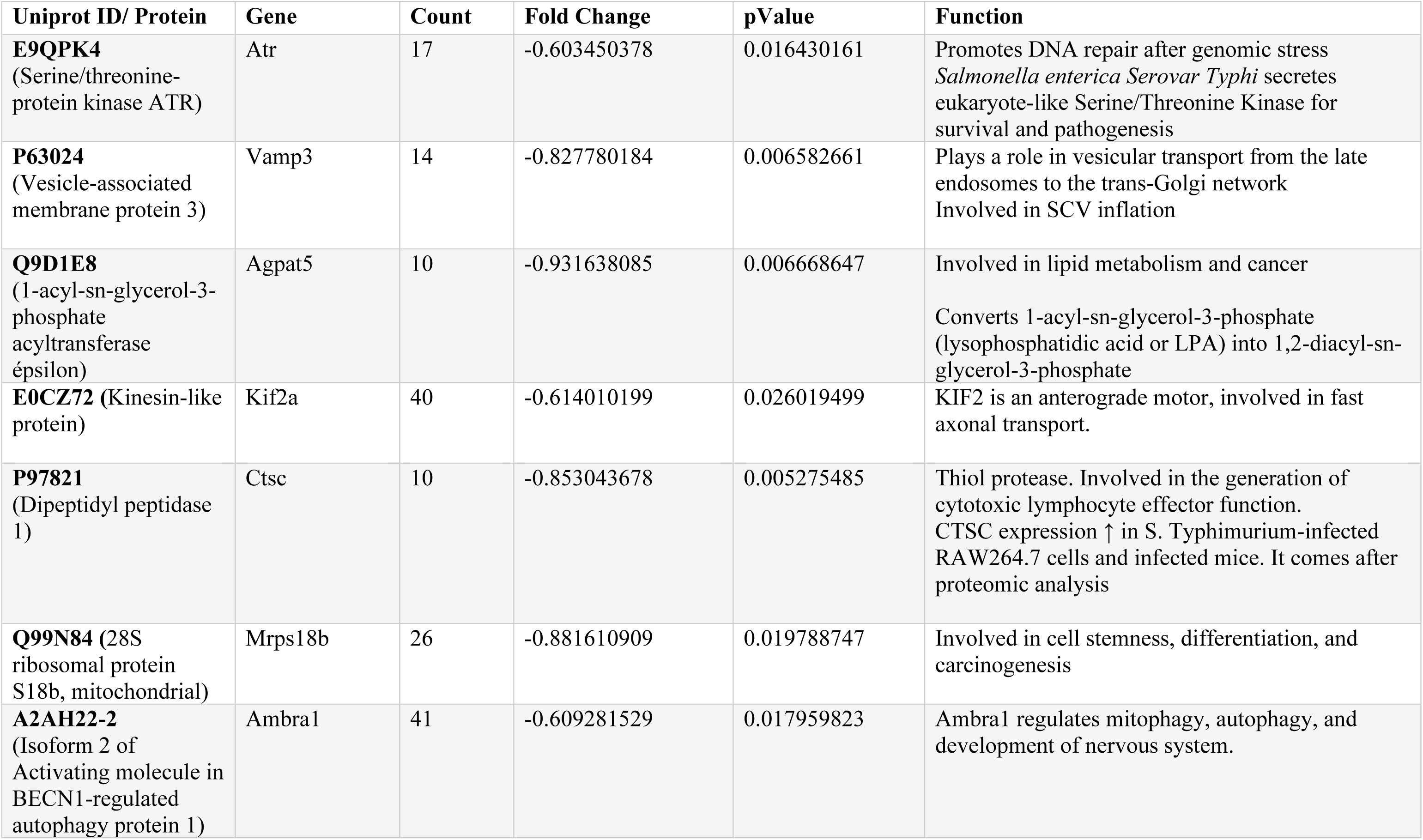
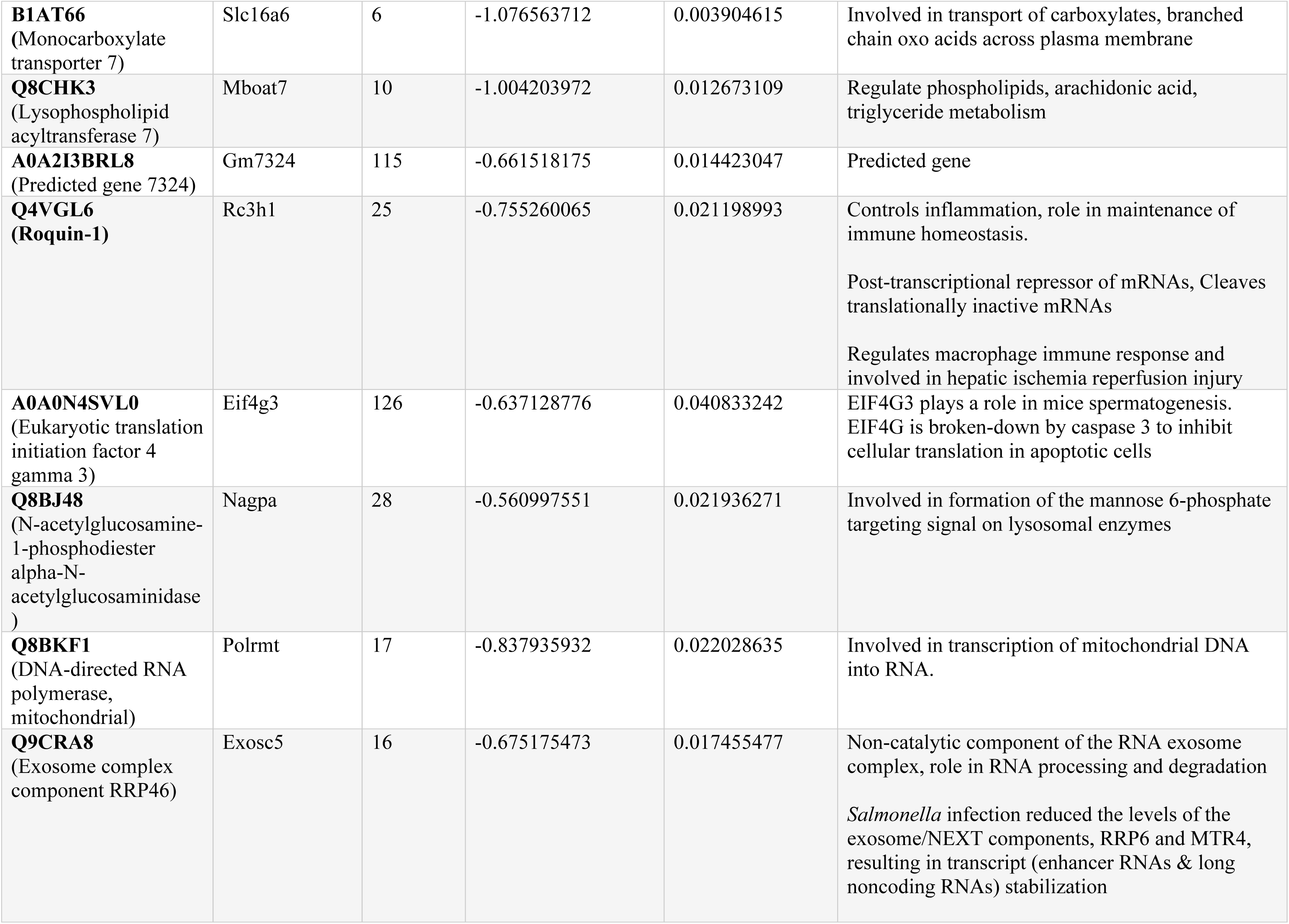
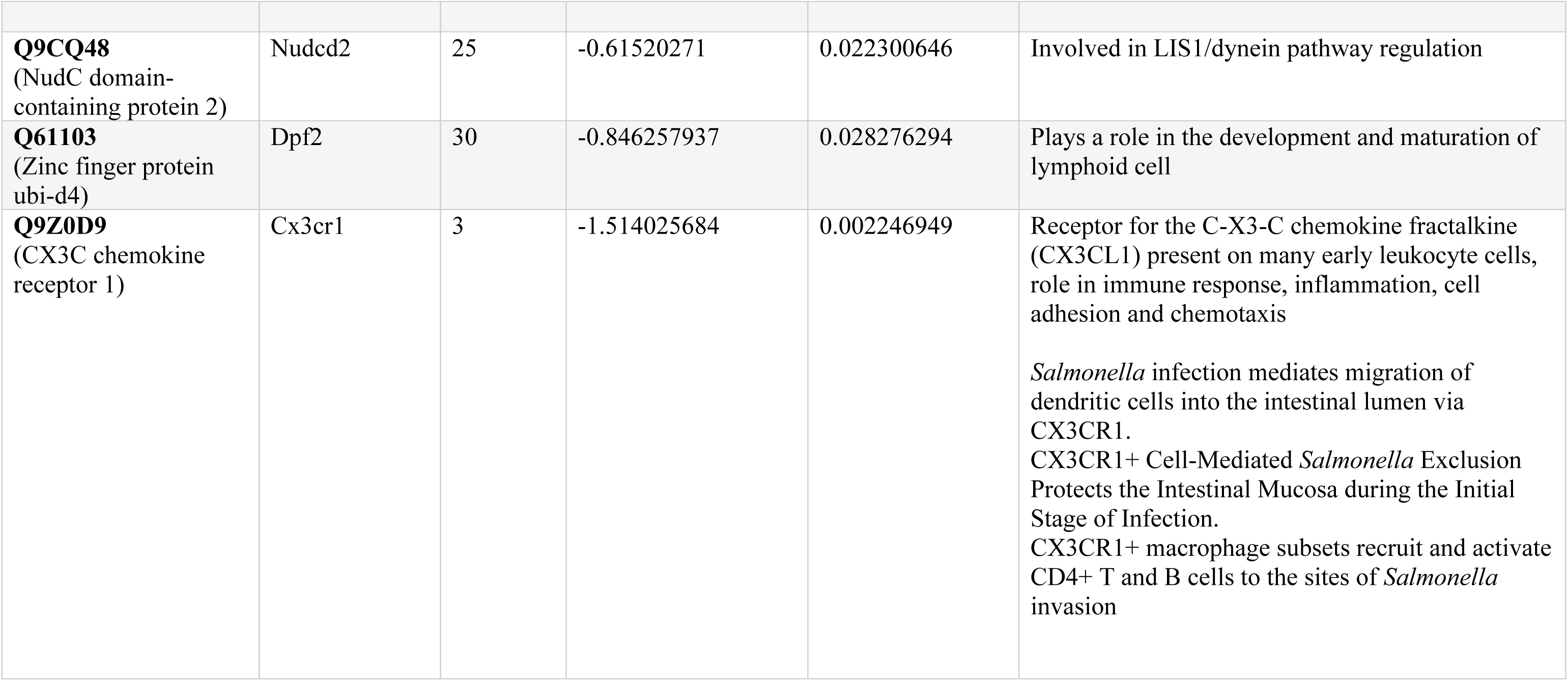
Functions of 18 downregulated proteins unique to E1-*SifA* conditions.

Additionally, E1 cells infected with *SifA* showed 3 upregulated and 17 downregulated proteins which were common to pairs 9 and 10 (Fig 7B), suggesting the contribution of ELMO1 and SifA in the functioning of these proteins. The upregulated proteins were associated with GO terms including protein localization, lipid metabolism, and neutrophil activation while the downregulated proteins included biological processes related to cell differentiation, lipid metabolism, and organelle organization among others. Interestingly, 11 upregulated and 2 downregulated proteins were common in Pair 7 and Pair 8 and were associated with biological processes related to mitochondrial fission and apoptosis, protein transport and secretion, and histone methylation.

## Discussion

Recent breakthroughs in omics technology identified several host signaling pathways following infection (54, 55). Here, we have performed liquid chromatography multinotch MS3-tandem mass tag (TMT) multiplexed proteomics to determine the global quantification of proteins regulated by ELMO1 in macrophages after *Salmonella* infection. We identified 7000 proteins and unique proteins exclusive to ELMO1-depleted cells after *Salmonella* (*SL*) infection or controlled by ELMO1 and *Salmonella* effector SifA. We noticed that ELMO1 regulates proteins with functions related to mitochondria, metabolism, vesicle transport, and the immune system. Our findings reveal that infection of macrophages with *Salmonella* profoundly alters cytosolic and mitochondrial metabolism, shifting it from <u>oxidative phosphorylation to glycolysis</u>, aligning with previous studies that highlight *Salmonella* stimulation of aerobic glycolysis in macrophages. This metabolic shift, accompanied by a reduced tricarboxylic acid (TCA) cycle and oxidative phosphorylation, facilitates *Salmonella* replication in host cells and mice (40, 41). Importantly, the impact of *Salmonella*-induced changes in the host metabolism, specifically glycolysis and oxidative phosphorylation is accentuated by the depletion of ELMO1, indicating that ELMO1 is required for mitigating the effect of *Salmonella* on the immunometabolism.

Mitochondria are important cell organelles with a role in inducing apoptosis and maintaining immune homeostasis. Mitochondria regulate ATP production to meet cellular demands, calcium signaling, and antimicrobial reactive oxygen species (ROS) production – factors critical for eliciting an immune response to pathogenic infections (56). Therefore, pathogenic bacteria including *Salmonella* manipulate mitochondrial pathways to facilitate their growth. Our findings reveal a significant decrease in mitochondria respiration and ATP production during *Salmonella* infection. We also demonstrated that the absence of ELMO1 after *Salmonella* infection reduced ATP production and mitochondrial respiration. It needs further investigation to know the changes after *Salmonella* infection are all by ELMO1-dependent or by ELMO1-independent pathways.

Within the identified proteins, we observed a significant number of host proteins linked with mitochondrial function as listed in Table 2-4. Besides the proteins that we validated, we noticed that the mitochondrial ribosomal proteins MRPL12, 43, and 49 are downregulated in E1 *SL* compared to C1 *SL*. Mitochondrial ribosomes (mitoribosomes) perform protein synthesis inside mitochondria, the organelles responsible for energy conversion and adenosine triphosphate production in eukaryotic cells. This provides an additional indication that ELMO1 has a crucial role in controlling the overall functions of mitochondria. Dnm1l or DRP1 is the most upregulated (4-fold) protein in the ELMO1-depleted macrophages after *Salmonella* infection as observed from the unique proteins in Table 2. We further validated that DRP1 is regulated translationally and increased in ELMO1-depleted macrophage cell line as well as in ELMO1 KO mice when we used colonic tissue of mice infected with *SL*.

DRP1 is involved in mitochondrial fission (47, 48). The coordinated balance between mitochondrial fusion and fission, known as mitochondrial dynamics, regulates mitochondrial morphology and function. Mitochondrial fission has primarily been associated with pro-inflammatory responses and metabolic adaptation, so it can be considered combative strategy immune cells utilize. In contrast, mitochondrial fusion has a more protective role in limiting cell death under conditions of nutrient starvation(57). Alterations in mitochondrial dynamics are associated with cancer development, and progression of cardiovascular and neurodegenerative conditions (58–61). Emerging studies have shown that mitochondrial functions are involved in gastro-intestinal diseases including in the inflammatory bowel diseases (62). Mitochondrial dynamics also influence host-pathogen interaction and host defences and favor pathogen persistence, replication, and/or survival, depending on the nature of the pathogen and cellular context, as shown in *Listeria*(63), *Legionella* (64), *Helicobacter pylori* VacA (65) and *Shigella* (66). The mitochondrial pathway can activate pattern-recognition receptor to initiate immune responses to clear the pathogens. For example, TLR signalling triggers major changes in metabolic pathways in murine macrophages that connect TLR with mitochondrial function (67).

The upregulation of proteins involved in mitochondrial fission and biogenesis in ELMO1-depleted macrophages after *Salmonella* infection suggests a putative role for ELMO1 in controlling mitophagy and other apoptotic pathways critical for mitochondrial health. Interestingly, the proteins differentially controlled by ELMO1-SifA also belong to mitochondrial pathways and apoptosis (Nudt9 and Mrps18b); vesicle-mediated transport (SNX5); lipid metabolism, and regulation of the immune system. The link of these cellular signaling with *Salmonella* effector SifA will open new directions in host-microbe interaction. Our findings indicated that *Salmonella*, and probably other bacteria, reprogram the host metabolism and regulatory mechanisms inside immune cells to take the best advantage of metabolites present inside the hosts and these mechanisms are modulated by the host microbial sensor ELMO1.

## Abbreviations

AGC: Automatic gain control
ANOVA: Analysis of variance
ATP: Adenosine triphosphate
BAI1: Brain Angiogenesis Inhibitor1
BOLA1: BolA Family Member 1
CID: collision-induced dissociation
DAVID: Database for Annotation, Visualization and Integrated Discovery
DEP: differentially expressed protein
DMEM: Dulbecco’s Modified Eagle Medium
DOI: Digital Object Identifier
DRP1: Dynamin-related protein 1
ECAR: extracellular acidification rate
ELMO1: Engulfment and cell motility protein 1
ER: endoplasmic reticulum
FCCP: Carbonyl cyanide-p-trifluoromethoxyphenylhydrazone
GEF: guanine nucleotide exchange factor
GO: Gene Ontology
HCD: High Energy Collisional Dissociation
HEPES: (4-(2-hydroxyethyl)-1-piperazineethanesulfonic acid)
HRP: Horseradish Peroxidase
IAA: Iodoacetamide
IACUC: Institutional Animal Care and Use Committee
MS: Mass spectrometry
Myl6: myosin light chain peptide 6
NFkB: Nuclear factor kappa-light-chain-enhancer of activated B cells
OCR: oxygen consumption rate
OXPHOS: oxidative phosphorylation
PAMPs: pathogen-associated molecular patterns
PBS: Phosphate-buffered saline
PPI: protein-protein interaction
PRRs: Pattern Recognition Receptors
PVDF: polyvinylidene difluoride
RIG-1: Retinoic acid-inducible gene I
RIPA: Radioimmunoprecipitation assay buffer
ROS: reactive oxygen species
SCVs: Salmonella-containing vacuoles
SD: standard deviation
SDS/PAGE: Sodium dodecyl-sulfate polyacrylamide gel electrophoresis
SEM: standard error of the mean
SL: Salmonella
SNX5: Sorting Nexin 5
SPS: synchronous precursor selection
TCA: tricarboxylic acid cycle
TEAB: Triethylammonium bicarbonate
TFA: Trifluoroacetic acid
TGEb: transforming growth factor beta receptor signalling pathway
TNF-α: Tumor necrosis factor
TMT: Tandem Mass Tags
WT: Wild-type.

## Acknowledgments

We appreciate the technical support from Shanaan Mujuru and Samantha Graham, the student interns of the Das lab. Several illustrations present in the manuscript are designed using www.biorender.com.

## Author Contribution

SD and DJG designed and planned the research; SCA, DM, ST, SRI, and IMS performed research; SD, IMS, and DJG, contributed new reagents/analytical tools; SCA, DM, ST, SRI, and IMS analyzed data; SD and DJG supervised the research. All authors wrote their respective parts and coordinated with SD for the manuscript draft. All authors edited and agreed on the final version of the manuscript.

## Data Sharing plan

All proteomics data has been submitted and all other raw data is accessible from the Cloud.

## Supplementary Figure Legends

**Figure S1.**
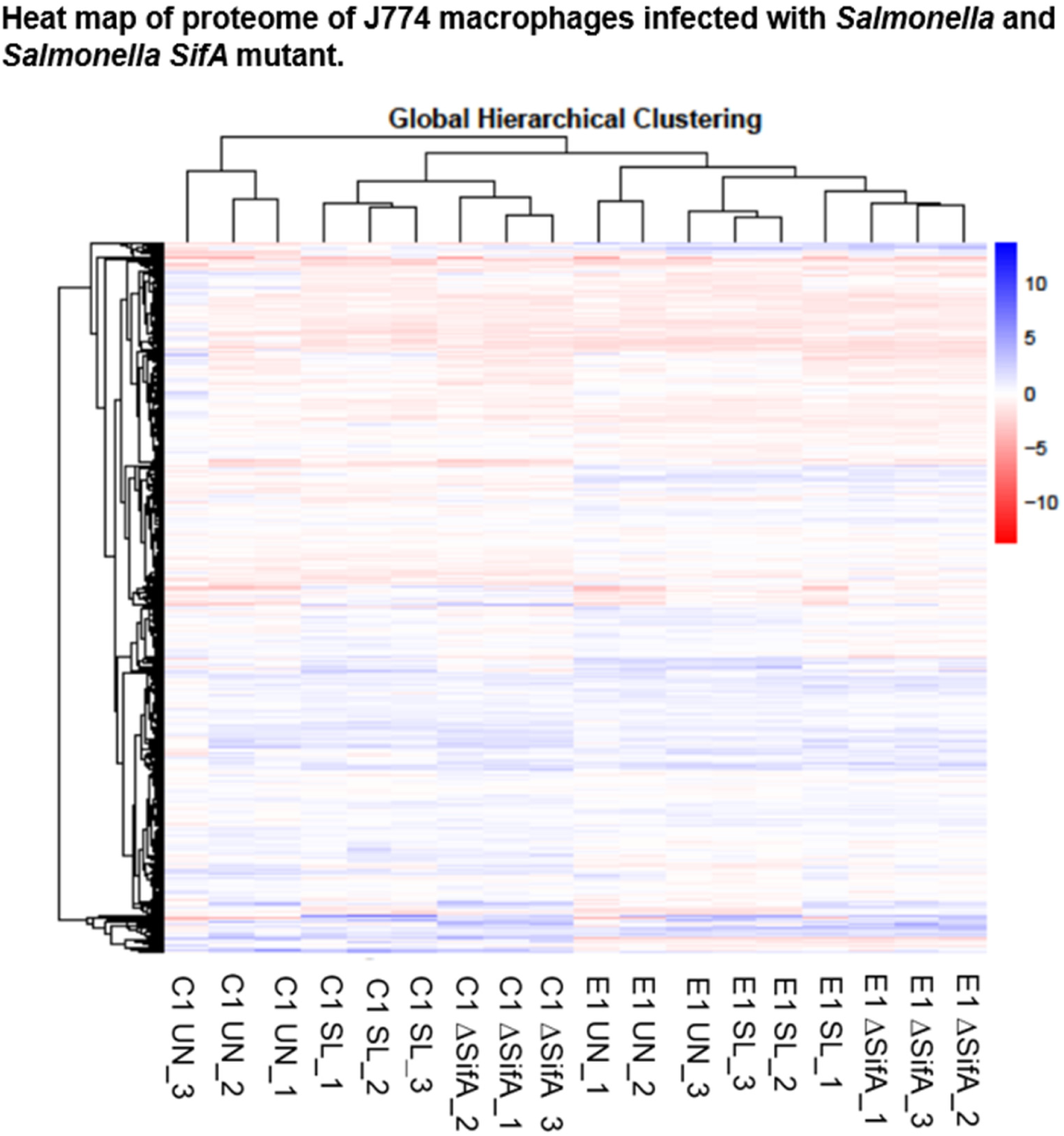
A heatmap showing the total number of differentially expressed hits identified by mass spectrometry (blue color is used for upregulated targets and red color for downregulated targets). Different sample groups (in columns) were arranged using a clustering algorithm. The box at the bottom shows the total number of peptide spectral matches (200,148) and protein IDs (7777) identified.

**Figure S2.**
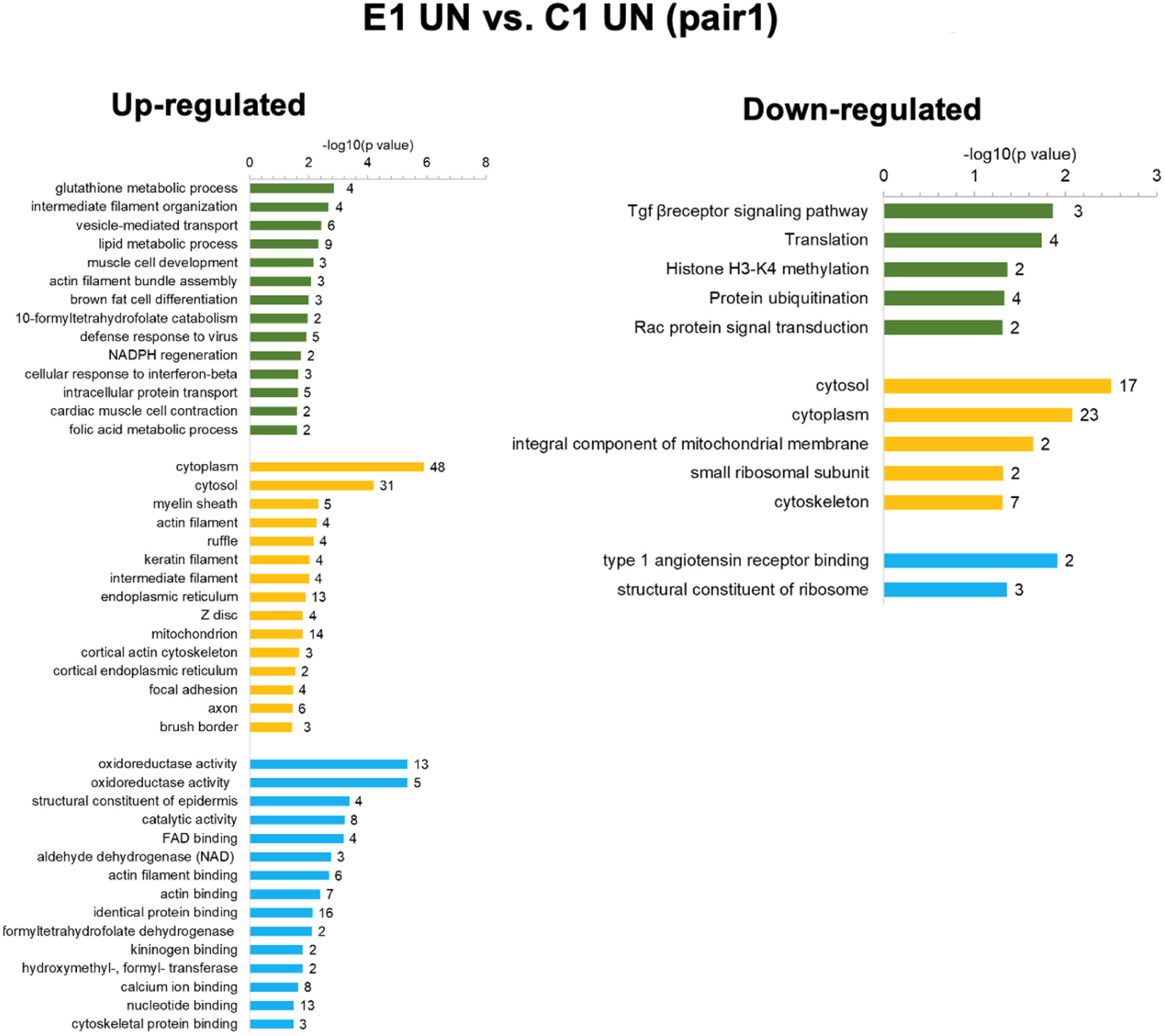
Representation of GO enrichment and DAVID Functional Annotations for analysis of 74 proteins upregulated proteins and 40 downregulated proteins in E1 UN vs C1 UN. Green Bars represent biological functions, yellow Bars represent cellular components, and blue bars represent molecular functions. The left panel represents upregulated proteins and the right panel represents downregulated ones.

**Figure S3.**
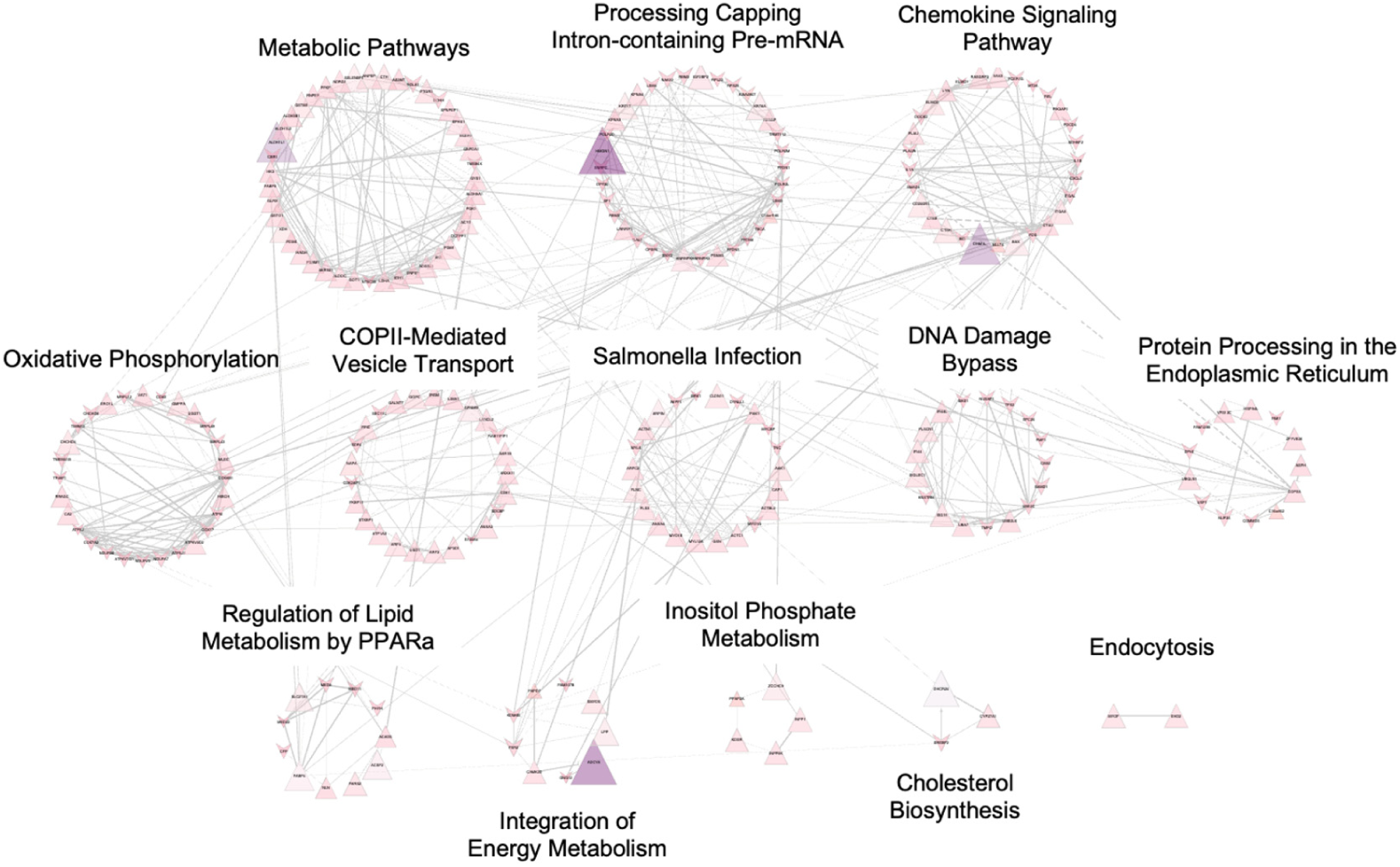
Protein-protein interaction (PPI) has been shown using STRING software to indicate pathways involved with ELMO1 following *Salmonella* infection.

**Figure S4.**
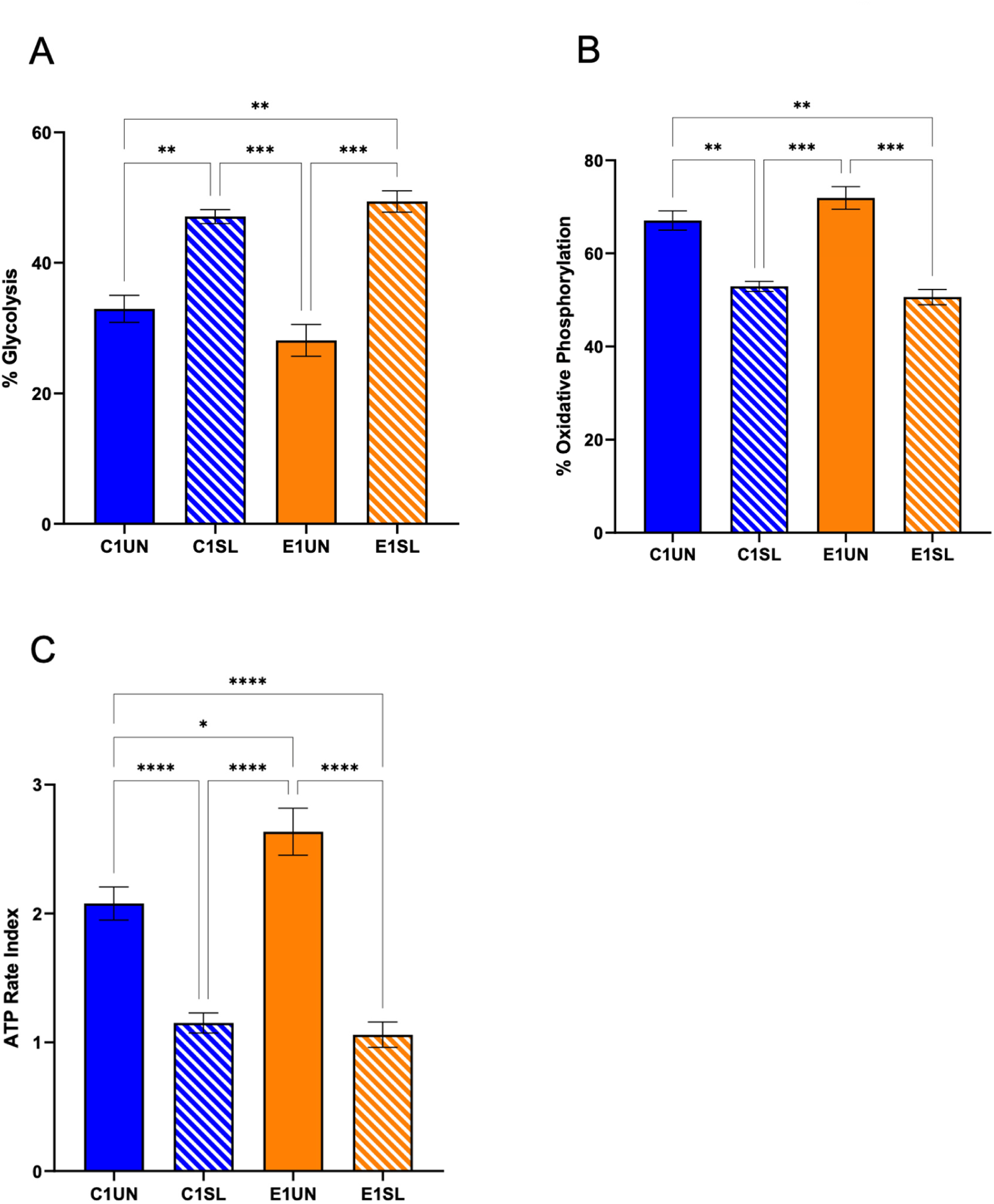
Quantification of the percentage of Glycolysis (A), percentage of Oxidative Phosphorylation (B), and ATP Rate Index (C) in C1 UN, E1 UN, and C1 and E1 following 6 h of infection with WT *SL* as measured by the instrument Agilent Seahorse XF HS Mini Analyzer. Ordinary one-way ANOVA was used for statistical analysis

**Figure S5.**
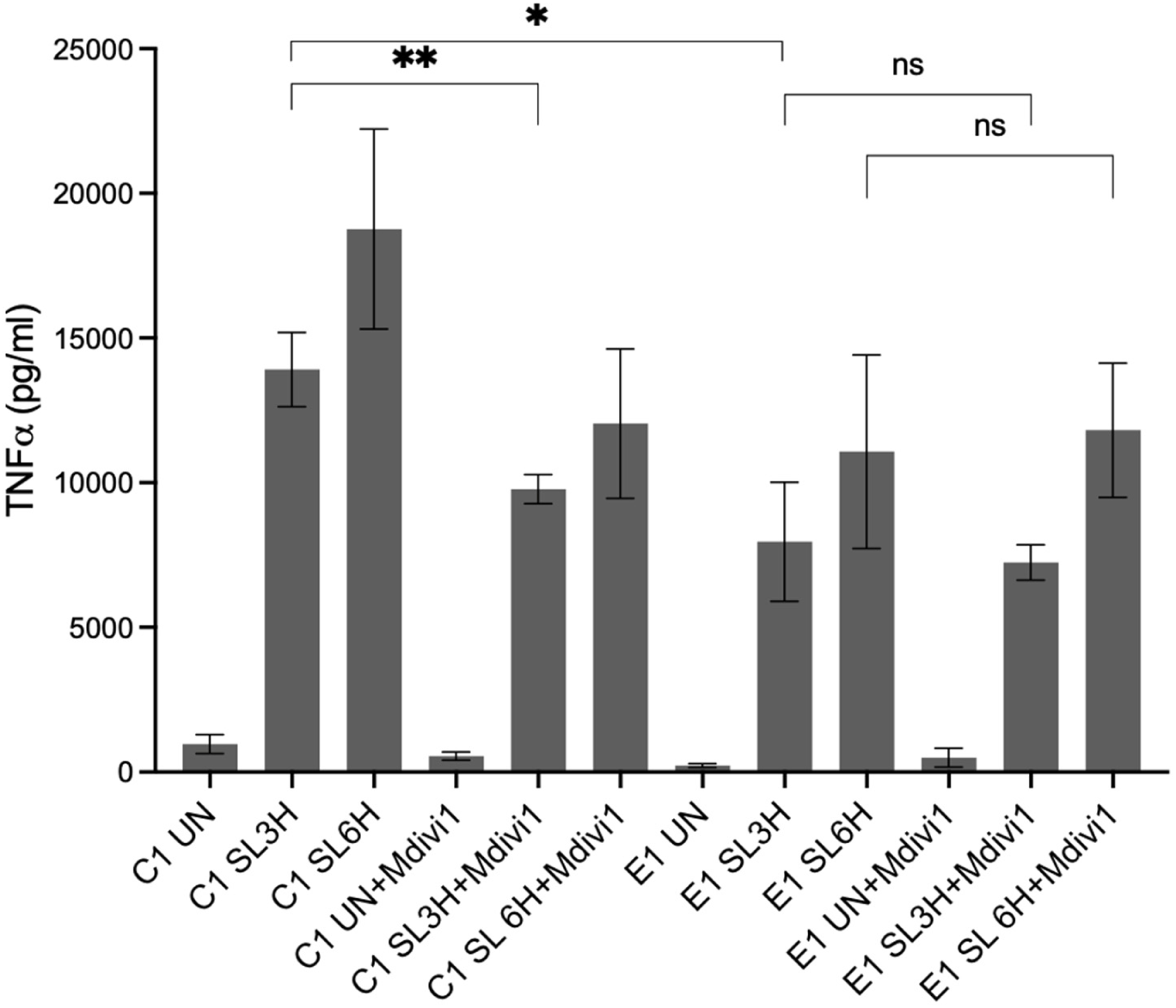
TNF-α cytokine levels in the supernatant from C1 and E1 cells after 3h and 6h infection with *SL* in the presence of DRP1 inhibitor mDivi-1, measured by ELISA. Data represents as the mean ± SEM of two separate experiments each including four biological replicates. * Indicates p ≤ 0.05, ** indicates p ≤ 0.01, as assayed by two-tailed Student’s *t*-test.

## References

1. D. Becker et al., Robust Salmonella metabolism limits possibilities for new antimicrobials. Nature 440, 303–307 (2006).

2. J. Noster et al., Proteomics of intracellular Salmonella enterica reveals roles of Salmonella pathogenicity island 2 in metabolism and antioxidant defense. PLoS Pathog 15, e1007741 (2019).

3. L. Shi et al., Proteomic analysis of Salmonella enterica serovar typhimurium isolated from RAW 264.7 macrophages: identification of a novel protein that contributes to the replication of serovar typhimurium inside macrophages. J Biol Chem 281, 29131–29140 (2006).

4. B. Steeb et al., Parallel exploitation of diverse host nutrients enhances Salmonella virulence. PLoS Pathog 9, e1003301 (2013).

5. L. Shi et al., Proteomic investigation of the time course responses of RAW 264.7 macrophages to infection with Salmonella enterica. Infect Immun 77, 3227–3233 (2009).

6. M. Hahn et al., SIK2 orchestrates actin-dependent host response upon Salmonella infection. Proc Natl Acad Sci U S A 118 (2021).

7. R. G. Jenner, R. A. Young, Insights into host responses against pathogens from transcriptional profiling. Nat Rev Microbiol 3, 281–294 (2005).

8. L. Qi et al., Quantitative proteomic analysis of host epithelial cells infected by Salmonella enterica serovar Typhimurium. Proteomics 17 (2017).

9. S. P. Salcedo, D. W. Holden, SseG, a virulence protein that targets Salmonella to the Golgi network. EMBO J 22, 5003–5014 (2003).

10. M. W. Vogels et al., Quantitative proteomic identification of host factors involved in the Salmonella typhimurium infection cycle. Proteomics 11, 4477–4491 (2011).

11. J. C. Santos et al., The COPII complex and lysosomal VAMP7 determine intracellular Salmonella localization and growth. Cell Microbiol 17, 1699–1720 (2015).

12. A. Thompson et al., Tandem mass tags: a novel quantification strategy for comparative analysis of complex protein mixtures by MS/MS. Anal Chem 75, 1895–1904 (2003).

13. M. Fukata, A. S. Vamadevan, M. T. Abreu, Toll-like receptors (TLRs) and Nod-like receptors (NLRs) in inflammatory disorders. Seminars in immunology 21, 242–253 (2009).

14. A. Pluddemann, S. Mukhopadhyay, S. Gordon, Innate immunity to intracellular pathogens: macrophage receptors and responses to microbial entry. Immunological reviews 240, 11–24 (2011).

15. K. Newton, V. M. Dixit, Signaling in innate immunity and inflammation. Cold Spring Harbor perspectives in biology 4 (2012).

16. D. Park et al., BAI1 is an engulfment receptor for apoptotic cells upstream of the ELMO/Dock180/Rac module. Nature 450, 430–434 (2007).

17. B. A. McCormick, ELMO1: More Than Just a Director of Phagocytosis. Cell Mol Gastroenterol Hepatol 1, 262–263 (2015).

18. S. Das et al., ELMO1 has an essential role in the internalization of Typhimurium into enteric macrophages that impacts disease outcome. Cell Mol Gastroenterol Hepatol 1, 311–324 (2015).

19. I. M. Sayed et al., Host engulfment pathway controls inflammation in inflammatory bowel disease. FEBS J 10.1111/febs.15236 (2020).

20. S. Tocci, S. R. Ibeawuchi, S. Das, I. M. Sayed, Role of ELMO1 in inflammation and cancer-clinical implications. Cell Oncol (Dordr) 45, 505–525 (2022).

21. A. Sarkar et al., ELMO1 Regulates Autophagy Induction and Bacterial Clearance During Enteric Infection. J Infect Dis 216, 1655–1666 (2017).

22. A. Sharma et al., The crosstalk between microbial sensors ELMO1 and NOD2 shape intestinal immune responses. Virulence 14, 2171690 (2023).

23. S. C. Achi, S. Karimilangi, D. Lie, I. M. Sayed, S. Das, The WxxxE proteins in microbial pathogenesis. Crit Rev Microbiol 49, 197–213 (2023).

24. I. M. Sayed et al., The interaction of enteric bacterial effectors with the host engulfment pathway control innate immune responses. Gut Microbes 13, 1991776 (2021).

25. J. D. Lapek Jr et al., Defining host responses during systemic bacterial infection through construction of a murine organ proteome atlas. Cell systems 6, 579–592. e574 (2018).

26. A. Thompson et al., Tandem mass tags: a novel quantification strategy for comparative analysis of complex protein mixtures by MS/MS. Analytical chemistry 75, 1895–1904 (2003).

27. I. H. Wierzbicki et al., Group A streptococcal S protein utilizes red blood cells as immune camouflage and is a critical determinant for immune evasion. Cell reports 29, 2979–2989. e2915 (2019).

28. J. K. Eng, A. L. McCormack, J. R. Yates, An approach to correlate tandem mass spectral data of peptides with amino acid sequences in a protein database. Journal of the american society for mass spectrometry 5, 976–989 (1994).

29. J. E. Elias, S. P. Gygi, Target-decoy search strategy for increased confidence in large-scale protein identifications by mass spectrometry. Nature methods 4, 207–214 (2007).

30. Y. Xiao et al., A novel significance score for gene selection and ranking. Bioinformatics 30, 801–807 (2014).

31. B. T. Sherman et al., DAVID: a web server for functional enrichment analysis and functional annotation of gene lists (2021 update). Nucleic Acids Res 10 (2022).

32. D. W. Huang, B. T. Sherman, R. A. Lempicki, Systematic and integrative analysis of large gene lists using DAVID bioinformatics resources. Nature protocols 4, 44–57 (2009).

33. Z. Xie et al., Gene set knowledge discovery with enrichr. Current protocols 1, e90 (2021).

34. M. V. Kuleshov et al., Enrichr: a comprehensive gene set enrichment analysis web server 2016 update. Nucleic acids research 44, W90–W97 (2016).

35. E. Y. Chen et al., Enrichr: interactive and collaborative HTML5 gene list enrichment analysis tool. BMC bioinformatics 14, 1–14 (2013).

36. M. Lu et al., PH domain of ELMO functions in trans to regulate Rac activation via Dock180. Nat Struct Mol Biol 11, 756–762 (2004).

37. S. Das et al., Brain angiogenesis inhibitor 1 (BAI1) is a pattern recognition receptor that mediates macrophage binding and engulfment of Gram-negative bacteria. Proc Natl Acad Sci U S A 108, 2136–2141 (2011).

38. Y. Handa et al., Shigella IpgB1 promotes bacterial entry through the ELMO-Dock180 machinery. Nat Cell Biol 9, 121–128 (2007).

39. S. D. Bowden, G. Rowley, J. C. Hinton, A. Thompson, Glucose and glycolysis are required for the successful infection of macrophages and mice by Salmonella enterica serovar typhimurium. Infect Immun 77, 3117–3126 (2009).

40. L. Jiang et al., Salmonella Typhimurium reprograms macrophage metabolism via T3SS effector SopE2 to promote intracellular replication and virulence. Nat Commun 12, 879 (2021).

41. J. Wang et al., Salmonella enterica Serovar Typhi Induces Host Metabolic Reprogramming to Increase Glucose Availability for Intracellular Replication. Int J Mol Sci 22 (2021).

42. P. Chaukimath, G. Frankel, S. S. Visweswariah, The metabolic impact of bacterial infection in the gut. FEBS J 290, 3928–3945 (2023).

43. Y. Wang et al., Host metabolic shift during systemic Salmonella infection revealed by comparative proteomics. Emerg Microbes Infect 10, 1849–1861 (2021).

44. T. Liwinski, D. Zheng, E. Elinav, The microbiome and cytosolic innate immune receptors. Immunological reviews 297, 207–224 (2020).

45. M. A. Uzarska et al., Mitochondrial Bol1 and Bol3 function as assembly factors for specific iron-sulfur proteins. Elife 5 (2016).

46. P. Willems et al., BOLA1 is an aerobic protein that prevents mitochondrial morphology changes induced by glutathione depletion. Antioxid Redox Signal 18, 129–138 (2013).

47. T. B. Fonseca, A. Sanchez-Guerrero, I. Milosevic, N. Raimundo, Mitochondrial fission requires DRP1 but not dynamins. Nature 570, E34–E42 (2019).

48. C. Hu, Y. Huang, L. Li, Drp1-Dependent Mitochondrial Fission Plays Critical Roles in Physiological and Pathological Progresses in Mammals. Int J Mol Sci 18 (2017).

49. R. Kapetanovic et al., Lipopolysaccharide promotes Drp1-dependent mitochondrial fission and associated inflammatory responses in macrophages. Immunol Cell Biol 98, 528–539 (2020).

50. V. Tiku, M. W. Tan, I. Dikic, Mitochondrial Functions in Infection and Immunity: (Trends in Cell Biology 30, 263-275, 2020). Trends Cell Biol 30, 748 (2020).

51. A. Cassidy-Stone et al., Chemical inhibition of the mitochondrial division dynamin reveals its role in Bax/Bak-dependent mitochondrial outer membrane permeabilization. Dev Cell 14, 193–204 (2008).

52. Q. Sun et al., Structural and functional insights into sorting nexin 5/6 interaction with bacterial effector IncE. Signal Transduct Target Ther 2, 17030 (2017).

53. T. Matsumiya, D. M. Stafforini, Function and regulation of retinoic acid-inducible gene-I. Crit Rev Immunol 30, 489–513 (2010).

54. A. R. Al-Maleki, K. Braima, N. A. Rosli, Editorial: Integrated omics approaches in the understanding of host-pathogen interactions. Front Cell Infect Microbiol 13, 1215104 (2023).

55. M. M. Khan et al., Multi-Omics Strategies Uncover Host-Pathogen Interactions. ACS Infect Dis 5, 493–505 (2019).

56. V. Tiku, M. W. Tan, I. Dikic, Mitochondrial Functions in Infection and Immunity. Trends Cell Biol 30, 263–275 (2020).

57. S. F. Afroz et al., Mitochondrial dynamics in macrophages: divide to conquer or unite to survive? Biochem Soc Trans 51, 41–56 (2023).

58. W. Chen, H. Zhao, Y. Li, Mitochondrial dynamics in health and disease: mechanisms and potential targets. Signal Transduct Target Ther 8, 333 (2023).

59. G. W. Dorn, 2nd, Evolving Concepts of Mitochondrial Dynamics. Annu Rev Physiol 81, 1–17 (2019).

60. S. Kyriakoudi, A. Drousiotou, P. P. Petrou, When the Balance Tips: Dysregulation of Mitochondrial Dynamics as a Culprit in Disease. Int J Mol Sci 22 (2021).

61. Y. Ma, L. Wang, R. Jia, The role of mitochondrial dynamics in human cancers. Am J Cancer Res 10, 1278–1293 (2020).

62. G. T. Ho, A. L. Theiss, Mitochondria and Inflammatory Bowel Diseases: Toward a Stratified Therapeutic Intervention. Annu Rev Physiol 84, 435–459 (2022).

63. F. Stavru, F. Bouillaud, A. Sartori, D. Ricquier, P. Cossart, Listeria monocytogenes transiently alters mitochondrial dynamics during infection. Proc Natl Acad Sci U S A 108, 3612–3617 (2011).

64. P. Escoll et al., Legionella pneumophila Modulates Mitochondrial Dynamics to Trigger Metabolic Repurposing of Infected Macrophages. Cell Host Microbe 22, 302–316 e307 (2017).

65. A. Y. Seeger et al., Host cell sensing and restoration of mitochondrial function and metabolism within Helicobacter pylori VacA intoxicated cells. mBio 10.1128/mbio.02117-23, e0211723 (2023).

66. M. Lum, R. Morona, Dynamin-related protein Drp1 and mitochondria are important for Shigella flexneri infection. Int J Med Microbiol 304, 530–541 (2014).

67. A. P. West et al., TLR signalling augments macrophage bactericidal activity through mitochondrial ROS. Nature 472, 476–480 (2011).

